# Physiological and molecular responses to combinatorial iron and phosphate deficiencies in hexaploid wheat seedlings

**DOI:** 10.1101/2020.05.26.117101

**Authors:** Gazaldeep Kaur, Vishnu Shukla, Varsha Meena, Anil Kumar, Deepshikha Tyagi, Jagtar Singh, Pramod Kaitheri Kandoth, Shrikant Mantri, Hatem Rouached, Ajay Kumar Pandey

**Author notes:** Corresponding author* Dr. Ajay K Pandey, Scientist-F. National Agri-Food Biotechnology Institute (Department of Biotechnology), Sector 81, Knowledge City, Mohali-140306, Punjab, India. Both the authors contributed equally (Gazaldeep Kaur, Vishnu Shukla).

## Abstract

Iron (Fe) and phosphorus (P) are the essential mineral nutrient for plant growth and development. However, the molecular interaction of the Fe and P pathways in crops remained largely obscure. In this study, we provide a comprehensive physiological and molecular analysis of hexaploid wheat response to single (Fe, P) and its combinatorial deficiencies. Our data showed that inhibition of the primary root growth occurs in response to Fe deficiency; however, growth was rescued when combinatorial deficiencies occurred. Analysis of RNAseq revealed that distinct molecular rearrangements during combined deficiencies with predominance for genes related to metabolic pathways and secondary metabolite biosynthesis primarily include genes for UDP-glycosyltransferase, cytochrome-P450s, and glutathione metabolism. Interestingly, the Fe-responsive cis-regulatory elements in the roots in Fe stress conditions were enriched compared to the combined stress. Our metabolome data also revealed the accumulation of distinct metabolites such as amino-isobutyric acid, arabinonic acid, and aconitic acid in the combined stress environment. Overall, these results are essential in developing new strategies to improve the resilience of crops in limited nutrients.

**HIGHLIGHTS:**

- This study was performed to understand the molecular changes occurring during the interaction of Phosphorus (P) and Iron (Fe) in hexaploid wheat roots.
- P and Fe show cross-talk as Fe deficiency-induced phenotype that was restored by the withdrawal of P.
- A total of 2780 differentially expressed genes were identified in the roots with the combined –Fe–P deficiencies with predominance for UDP-glycosyltransferases, cytochrome-450 and glutathione-S-transferases transcripts.
- The metabolomic changes identified the importance of amino-isobutyric acid, arabinonic acid and aconitic acid during dual deficiency
- This work provides a comprehensive insight to understand the molecular rearrangements occurring in wheat roots during Fe and P interaction.

## 1. Introduction

Nutrient deficiencies in the rhizospheric environment of the plants severely reduce crop yields and negatively affect the worldwide food supply [1]. Iron (Fe) is an essential microelement for plant growth and development and is utilized in nearly every cellular process, including photosynthesis and respiration. In general, deficiency of Fe is considered as one of the most critical limitations in cereal crop production, especially in alkaline calcareous soil [2,3].To overcome Fe deficiency, non-graminaceous and graminaceous species rely on different strategies for transporting the available Fe. The strategy I is used by non-graminaceous plants wherein the lowering of rhizospheric pH is attained through excretion of protons with P-type ATPases coupled with Ferric reductase oxidase activity at the root plasma membrane, thereby causing reduction of Fe^3+^ to Fe^2+^, thereby making bioavailable to plant [4]. The uptake of Fe^2+^ is done by the divalent transporter Iron-regulated transporter (IRT1) [5,6]. Graminaceous plants rely on Strategy II, which uses chelation for Fe absorption, predominately through biosynthesis and secretion of phytosiderophores (PS), such as mugineic acids (MAs). These biochemicals make Fe-PS complexes that are subsequently transported into the roots by yellow stripe-like transporter proteins (YSL) [7–12]. The molecular players involved in the efflux of MA are now characterized in rice, barley and very recently identified in wheat [11,12]. Regulation of Fe acquisition, transport and mobilization mainly involve central transcription factors (TFs) belonging to the family of basic helix-loop-helix (bHLH), Fe□deficiency□responsive elements binding factors (IDEF1 and IDEF2) and IRO2/IRO3 [13–17].

Phosphorus (P) is an essential macronutrient for plants to complete their life cycle. The P deprivation typically results in root architecture changes to enhance its uptake and mobilization [18,19]. Under low P concentrations, the model plant *Arabidopsis* (Col-0) show reduced primary root elongation. This phenotype was proposed to be partly due to Fe toxicity at the root tip. Unlike *Arabidopsis*, most monocot plants, including maize, rice and wheat, showed either no reduction or slight elongation of primary roots under P deficiency [20–22]. In contrast, no Fe toxicity was reported in cereal crops grown under P deficient condition. Thus, how monocots (esp. cereal crops) coordinate P and Fe remains largely unknown. Based on the available information, the key components involved in adapting to P deficiency include the transcription factor (TF) PHOSPHATE STARVATION RESPONSE (*PHR1*); microRNA399 (miR399); PHOSPHATE1 (*PHO1*), SPX containing proteins, and ubiquitin E2-conjugase(*PHO2*) are functionally conserved in *Arabidopsis* (dicots), rice and wheat [23–25]. During the P deficiency, genes response is regulated by the TF (PHR1) though cis-elements PHR1 binding site (P1BS motif) present in their promoters. Overall, this suggests that roots undergo molecular reprogramming to cope with the nutrient stress.

When P and Fe are absorbed simultaneously, their availability is still affected by precipitation [26]. Multiple studies have provided preliminary insights into the interaction of P with the absorption of essential micronutrients like Fe [27,28]. Earlier studies have shown that P deficiency could promote iron content and storage in Arabidopsis tissue. This suggests that Fe is more accessible in the rhizosphere under P deficiency, which leads to reduced expression of Fe homeostasis-related genes [18,29,30]. Interestingly, genes such as ferritin and zinc-regulated, iron-regulated transporter-like proteins (ZIP) were speculated to be involved in enhancing Fe content in Arabidopsis [17,14,34]. The single nutrient deficiency (P and Fe) response has primarily been undertaken, but the simultaneous dual deficiency response remained limited only to Arabidopsis and rice [20]. In addition to this, the P deficiency also causes multiple biochemical and molecular changes that are also involved in Fe and Zinc (Zn) metabolism [32]. Genome-wide information was generated that revealed multiple protein-encoding genes from roots showing the overlap between Fe and P deficiency response [33]. Also, at the biochemical level, the dual deficiency also regulates the type of metabolites, including selective coumarins that are secreted by the plants [27]. Therefore, the detailed understanding between P and Fe homeostasis could provide a powerful tool to conduct molecular studies on nutrient signalling cross-talk for addressing the changing environmental scenarios for crop productivity [34].

Wheat is an important crop and is the major source of nutrition. Studying the molecular attributes of Fe and P cross-talk will help in designing a suitable model to optimize crop productivity during nutrient deficiency. In the current study, we performed a comprehensive analysis of physiological, transcriptional and metabolic changes during the combinatorial deficiencies of Fe and P (–Fe–P). An important role of genes involved in Fe uptake under the presence and absence of P was observed. Our work utilizes physiological and molecular evidences that identifies the distinct transcriptional and metabolic regulation in the roots of wheat during Fe and P interaction. Knowledge gained in this study expands our understanding of single and simultaneous nutrient stresses in cereal crops that are coordinated at the whole plant level.

## 2. Materials and Methods

### 2.1. Plant materials

Indian hexaploid wheat variety, ‘C-306’, a landrace genotype is known for its superior chapatti quality, was used for this study [35]. The ‘C-306’ seedlings were grown hydroponically under the Hoagland’s nutrient solution containing (L^-1^): 6 mM KNO_3_, 1mM MgSO_4_.7H_2_O, 2 mM Ca(NO_3_).4H_2_O, 200 µM KH_2_PO_4_, 20 µM Fe(III)EDTA, 0.25 mM H_3_BO_3_, 2 µM MnSO_4_.H_2_O, 2 µM ZnSO_4_.7H_2_O, 0.5 µM CuSO_4_.5H_2_O, 0.5 µM Na_2_.MoO_4_ and 0.05 mM KCL. After overnight stratification at 4 °C, wheat seeds were germinated for 5 days in moist (MiliQ water) filter paper. Once the endosperm starts browning, it is removed from the developing seedlings. Seedlings were then transferred to PhytaBox™ and grown in the nutrient solution described above. After 7 days, nutrient solutions were replaced on the basis of different treatments. For +Fe–P treatment, 20 µM KH_2_PO_4_ was used that was shown earlier for P deficiency [36]. For – Fe+P treatment, 2 µM Fe (III) EDTA was used for Fe deficiency [12]. While for –Fe–P treatment, 20 µM KH_2_PO_4_ and 2 µM Fe (III) EDTA were used. Control plants (+Fe+P) were grown in the above-listed concentrations of Hoagland’s solution. Plants were grown in the described medium for 20 days in a growth chamber set at 20□±□1□°C, 50–70% relative humidity and photon rate of 300□μmol quanta m^−2^ s^−1^ with 16□h□day/8□h night cycle. The whole set of experiments was repeated four times to examine biological variation. For sampling, all the roots and shoots were collected post 20 days after treatment (DAT; at the Zadoks growth scale, Z15) and at two hours after the onset of light. Samples were snap-frozen in liquid nitrogen and stored at -80°C. To characterize the primary root and 1^st^ order lateral root, eight individual plants of each of the three replicates were moved onto 150mm wide petri-plates filled with distilled water and examined manually.

### 2.2 RNASeq experiment design and sequencing

Total RNA extraction was done using twenty-day old root tissue samples for the four conditions (+Fe+P, +Fe–P, –Fe+P and –Fe–P). In total, three biological replicates derived from at least three independent events were sampled for RNA extraction. Each biological replicate consisted of 12-15 seedlings per treatment. Samples were collected at the same time, snap-frozen in liquid nitrogen and stored at -80°C. The RNA extraction and Illumina sequencing were performed with three replicates for –P and three replicates for –Fe–P with their respective controls. For Fe deficiency, previously published RNAseq datasets were used [12]. All the treatments to the seedlings were carried out during the same time, although the analysis was carried earlier from –Fe+P RNA samples. RNA extraction for library construction from the control and treated root samples were performed as reported previously [12]. Briefly, sequence libraries were prepared from high quality, quality-control passed RNA samples using Illumina

TruSeq mRNA library prep kit as per the instructions (Illumina Inc., USA). The reads were sequenced using 2 × 150 bp chemistry on NextSeq 500 and NovaSeq 6000. Only one biological replicate for –P did not generate the desired output data size; therefore, only two replicate data sets were used for subsequent analysis. These replicates were validated for their conserved P deficiency response as noted previously [24] before proceeding for study.

### 2.3. Sample clustering and differential expression analysis

Paired-end reads were quality trimmed and adapter filtered using Trimmomatic v0.35 to retain only good quality reads (QV>20). The clean raw reads were quantified for expression by pseudoalignment against wheat transcriptome (ensembl release 46; ftp://ftp.ensemblgenomes.org/pub/plants/release-46/fasta/triticum_aestivum/ /IWGSC RefSeq v1.0) using Kallisto v0.44.0 [37] and the option --rf-stranded for stranded samples. DESeq2 R package [38] was used for differential expression analysis. Raw count values from Kallisto were summarized from transcript to gene-level abundances and imported for use with DESeq2 using tximport package [39]. Principal Component Analysis (PCA) and clustered heatmap analysis was performed based on VST transformed counts for the top 500 genes showing highest variation in expression to observe clustering across replicates and conditions. The representatives from batches were taken together to build a DESeq2 model to obtain normalised counts, which were transformed using VST mode and corrected for the associated batch effect using remove Batch Effect function from limma package [40]. The ggplot2 [41] and pheatmap package were used to design the PCA plot and clustered heatmap, respectively.

The DESeq() function was used for differential expression analysis to calculate a relative expression for pairwise comparisons among conditions. Log_2_ Fold Changes (Log2FC) were obtained for the pairwise comparisons for each of the three deficiency conditions and the control (+Fe+P) from the respective batch group only to further avoid any variation due to batch effects. The relative expression ratios were shrunk using apeglm package [42] to adjust the Log2FC of genes with extremely low counts. The percentage for genes with similar expression changes (overlapping DEG percentage) under two different conditions with respect to control was calculated as follows: 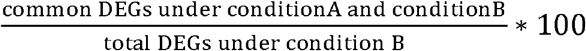

### 2.4 Functional enrichment analysis and annotation

KOBAS (KEGG Orthology-Based Annotation System) standalone tool was used to annotate wheat genes based on blast mapping against rice RefSeq and RAP-DB sequences, e-value <10^−5^. For pathway enrichment analysis, the “identify module” was used to shortlist the significantly overrepresented KEGG pathways for the respective deficiency conditions using Fisher’s exact test. FDR correction was performed using Benjamini and Hochberg method. Pathway enrichment results were depicted through bubble plots using ggplot2 package [41]. The size of the bubbles represents the number of genes altered in the respective pathways, rich factor on y-axis was calculated as a ratio of the number of genes altered to the total number of genes in a specific pathway. This means the greater the rich factor, the more intensive the enrichment in a pathway. Also, MapMan [43] was used to map the differentially expressed genes (DEGs) onto metabolic, regulatory and other biological categorical pathways. The mapping file was generated through Mercator [44] using the wheat transcriptome fasta file as an input. In addition, wheat RefSeq v1.1 annotation released by International Wheat Genome Sequencing Consortium (IWGSC) was also used (https://urgi.versailles.inra.fr/download/iwgsc).

### 2.5. Identification of cis-regulatory elements

To check the extent of Fe- or P-specific transcriptional regulation in single (Fe or P) and their combined deficiency, sequences for the 1500 bp upstream promoter region for each set of DEGs were downloaded from Ensembl Biomart. The promoter sequences were checked for the presence of 115 frequent cis-regulatory elements (freq-CREs) enriched in clusters for gold standard (GS) Fe responsive genes [14] and phosphate regulation specific CREs using an in-house perl script (https://github.com/Gazal90/FindMotifs/blob/master/motifs.pl). For validation and comparison, three sets of control groups with 100 promoters each were randomly shortlisted from genes that were not altered in response to Fe deficiency (−0.5 >Log2FC< 0.5).

### 2.6. Gas chromatography-mass spectrometry metabolite profiling

Extraction of total metabolites was performed similarly as previously described [45]. Wheat roots subjected to +Fe+P, +Fe–P, –P+Fe and –Fe–P were sampled at 20 days after deficiency in a triplicate manner and processed for metabolite extraction. The derivatized metabolites were analyzed with a GC instrument (Agilent technologies 7890, USA) coupled with mass spectrometry. The measurement from an injection volume of 1 µl was taken in the split-less mode in DB-5 column (30 m × 0.25 mm, 0.25 µm film thickness, Agilent) using helium as carrier gas. For analysis, qualitative analysis of chromatograms was performed in MassHunter Qualitative analysis Sp1 workstation (Agilent, USA). Identification and annotation of each compound was supervised manually using AMDIS software and NIST08 database (http://www.nist.gov/srd/mslist.html). Values were normalized to sample weight and internal control (sorbitol). Statistical analysis was performed as described earlier [46]. Log_2_ ratios of metabolite abundances in tested conditions were plotted against control conditions (+Fe+P). Delta method approximation was used to calculate standard errors (se) between test (T) and control (C) conditions, se log-ratio = 1/ln 2√[(SE_T_ /T)^2^ + (SE_C_ /C)^2^], where SE_T_ and SE_C_ are standard errors of average test and control metabolite abundances. For PCA and hierarchical clustering analysis, the clustv is (https://biit.cs.ut.ee/clustvis/) online program package with Euclidean distance as the similarity measure with complete linkage was used. A tab-delimited file of annotated metabolites with their corresponding log-transformed concentration values was used as input.

### 2.7. Quantitative real time-PCR analysis

Total RNA was isolated from the roots of the 20 DAT seedlings. A total of two µg of total RNA was used to prepare cDNA by using SuperScript III First-Strand Synthesis System (Invitrogen, USA). Samples were pre-treated with TURBO DNA-free kit to remove genomic contamination (Ambion, TX, USA). To perform quantitative RT-PCR (qRT-PCR) amplification was performed using conserved gene-specific primers capable of amplifying all the homoeologs (Table S1) along with internal control ARF (ADP-Ribosylation Factor) to normalize the expression data for each gene by using the Ct method [2^(−ΔΔCt)^] in the CFX96™ Real-Time PCR System (BioRadInc, USA). Three independent replicates with two-three technical replicates were performed for each sample [47].

### 2.8. Metal analysis in wheat tissues, SPAD measurements, Pi estimation, Perl staining and root Fe mobilization assay

Different metal analysis was performed using Inductive Coupled Plasma-Mass Spectrometry (ICP-MS, Agilent, USA). Metal analysis was performed as described previously [48,49]. Briefly, the mature seeds were grounded to a fine powder and subsequently subjected to the microwave-digested with HNO_3_ protocol (SuraPure™, Merck, USA). Respective metal standards were also prepared for analysis. The total chlorophyll content was expressed in terms of SPAD value. This was determined on the fully expanded youngest leaves (3-4 spots per leaf) of seedlings using a portable chlorophyll meter (Konica Minolta SPAD-502, Japan). Eight to ten seedlings were used for these measurements for each treatment. Experiments were repeated thrice with similar observations. Three independent replicates were performed for each 20 DAT sample. Pi concentration in wheat roots was measured by the molybdate-blue colorimetric method with a minimum of three biological replicates (containing n=8) [50]. For Perl staining, wheat roots were incubated with Perl’s Prussian blue (PPB) method consisting of equal amounts of premixed solution of 4% (v/v) HCL and 4% (w/v) K_4_Fe(CN)_6_.3H_2_O. Roots of eight to ten wheat seedlings from each experiment were taken and incubated in the mixture for 30 mins. Blue colour Fe-plaques were observed for the presence of Fe on the wheat roots seedlings, and representative images (five seedlings per treatment) were taken for respective treatments. To estimate the Fe remobilization ability of these roots, assays was done in the aerobic condition as described earlier in detail [12,51]. Based on these mobilization assays, Fe-solubilizing capacity was measured. Twenty-five wheat seedlings from each treatment were used for release in 60 ml of deionised water in the presence of Micropur (Katadyn, Switzerland). The final concentration of the released ferrous ion was estimated by measuring OD at 562 nm.

### 2.9 Statistical data analysis and data availability

To identify significant differentially expressed genes, a cut-off criterion of Log2FC > 1 in either direction, with an adjusted p-value (padj) of less than 0.05 was set. The padj values were obtained by using the Benjamini and Hochberg approach for controlling the false discovery rate. One-way analysis of variance was performed by using Origin^R^ software. The different letters indicate significant differences of means, and the bars indicate standard deviation (SD). The RNAseq data generated during this study have been deposited under the NCBI SRA database BioProjectID PRJNA667481 and PRJNA529036.

## 3. RESULTS

### 3.1. Phenotypic characterization of Fe- and P-deficient wheat

We compared wheat growth under four given conditions: nutrient-sufficient (+Fe+P, control), single nutrient deficiency (–Fe+P and +Fe–P), and combined nutrient deficiency (–Fe–P). In the presence of P, –Fe plants gradually displayed shorter roots compared to the control plants and plants grown in –P and –Fe–P conditions (Figure 1A & B). Post 20 DAT, the SPAD values (young leaves) were also calculated that measure the severity of the nutrient deficiency. The – Fe+P treated seedlings show the lowest SPAD (13.7±1.2^b^), suggesting Fe deficiency impact, whereas no marked symptoms were noted for the +Fe+P (34.9±3.7^a^). Seedlings under dual stress i.e. –Fe –P, also show a reduction in SPAD (18.0±2.2^c^), whereas P-deficient seedling, +Fe –P, show 24.3±2.6^d^ of SPAD value. In addition, the primary root length shows a slight increase during –P condition.

**Figure 1:**
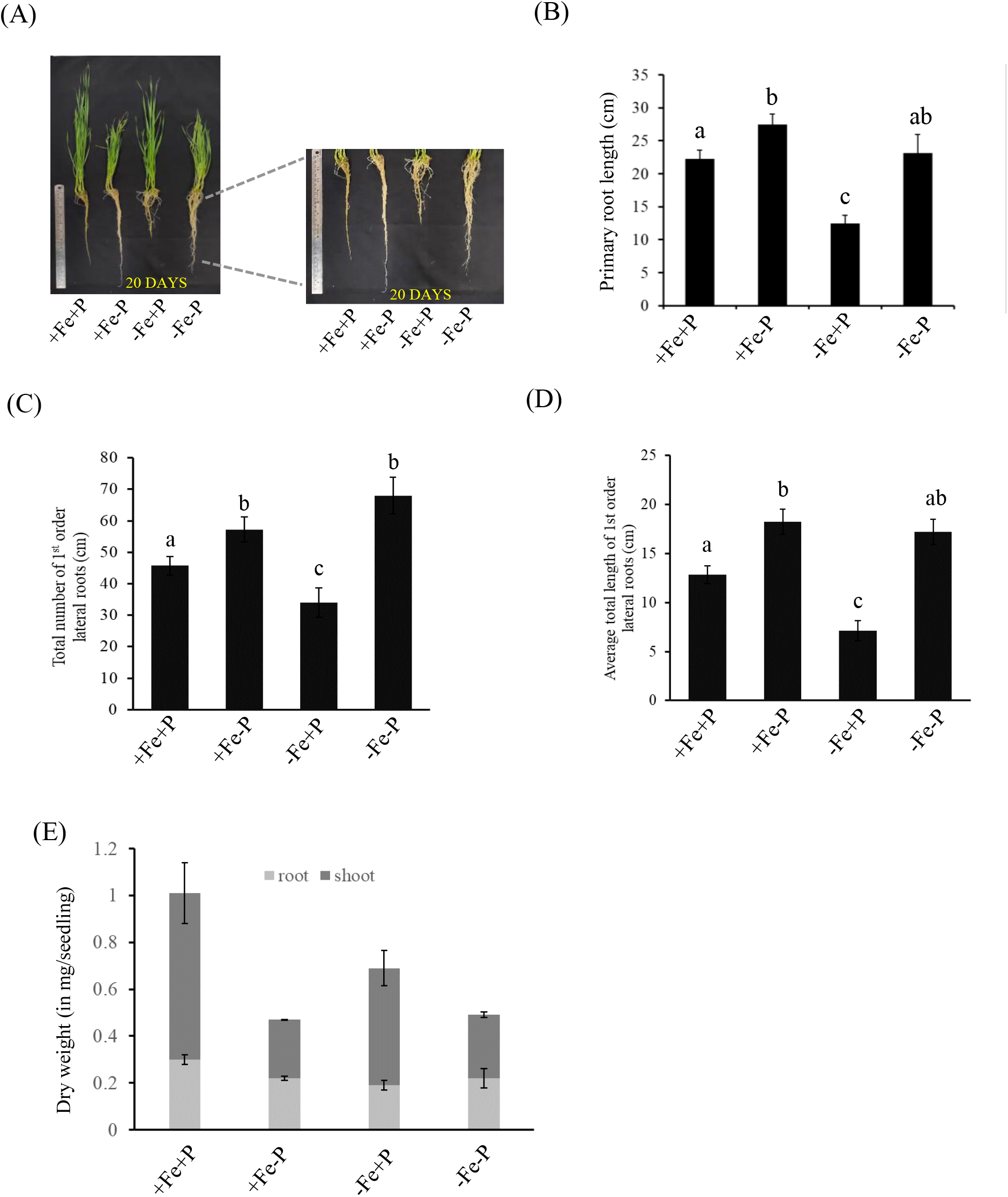
Effect of Fe and P on the growth of roots and shoots of wheat seedlings exposed to +Fe+P, +Fe–P, –Fe+P and –Fe–P after 20 days of treatment. (A) Experimental setup and the overall growth of wheat seedlings in the given conditions. Morphology of the wheat seedlings subjected to mentioned stress condition (+Fe+P, +Fe–P, –Fe+P and –Fe**–**P). Pictures were taken after 20 days of treatment. (B) Primary root length of wheat seedlings under-treated conditions. (C) The total number of 1^st^ order lateral roots and (D) Average total length of lateral roots. (E) Dry weight biomass of the seedlings (grams/seedling) of 20 days old seedlings under different mentioned conditions. One-way ANOVA with Tukey’s multiple comparison test means values was done. The different letters indicate a significant means difference, and the bars indicate SD (n=8 for B-E).

Interestingly, the –Fe–P condition led to the recovery of total primary root elongation, alleviating the negative effect of Fe deficiency (Figure 1B). A similar trend was also observed for the number and length of the 1^st^ order lateral roots, which shows a significant increase during –P and –Fe–P conditions relative to –Fe and control plants (Figure 1C and D). Examination of dry biomass of the seedling was assessed post 20 days of deficiency. The seedling biomass was higher in the control condition (∼1.01 grams per seedling) and lowest in –P and –Fe –P (∼0.47 and 0.49 grams per seedling) compared to –Fe (∼0.69 grams per seedling) (Figure 1E). Collectively our experiments suggest that Fe and P cross-talk could regulate the biomass allocation to roots and shoots, leading to the representative phenotype. This highlights that the deficiency of P on the wheat seedlings was more significant than those arising from the Fe deficiency.

### 3.2 Effect of deficiency conditions on nutrient content

Rhizospheric deficiency of Fe and P could affect the uptake of the metal ions in plants [52–55]. To check the influence of the single and dual deficiencies of P and Fe, we measured the concentration of Pi (inorganic phosphate), Zn and Manganese (Mn). During both –P or –Fe–P treatments reduced amount of Pi was measured. Interestingly, in 20 DAT roots, Pi content did not change under single –Fe+P deficiency (Figure 2A). Alternatively, significant increases in Fe accumulation occurred in the roots during –P (Table 1).

**Table 1:** Metal concentration in roots and shoots of wheat seedling subjected to –Fe, –P and –Fe–P stress. Different letters indicate significant changes within the treatments. One-way Annova and Duncan test (p>0.05) was performed for each within the treatments.

|  | Root |  |  | Shoot |  |  |
| --- | --- | --- | --- | --- | --- | --- |
| Treatment | Fe | Zn | Mn | Fe | Zn | Mn |
| (+Fe+P) | 160.91±4.66<sup>a</sup> | 27.1±0.88<sup>a</sup> | 59.53±2.87<sup>a</sup> | 86.87±4.19<sup>a</sup> | 24.43±1.76<sup>a</sup> | 21.52±0.94<sup>a</sup> |
| (+Fe-P) | 192.8±12.73<sup>b</sup> | 21.73±1.42<sup>b</sup> | 53.17±3.77<sup>b</sup> | 106.68±6.16<sup>b</sup> | 21.54±0.94<sup>a</sup> | 33.88±1.06<sup>b</sup> |
| (-Fe+P) | 115.44±25.37<sup>c</sup> | 29.25±2.21<sup>a</sup> | 28±1.55<sup>c</sup> | 40.13±2.31<sup>c</sup> | 32.36±2.48<sup>b</sup> | 26.83±0.66<sup>c</sup> |
| (-Fe-P) | 67.43±5.59<sup>d</sup> | 22.8±2.42<sup>b</sup> | 30.85±2.11<sup>c</sup> | 73.97±3.48<sup>d</sup> | 33.02±4.53<sup>b</sup> | 32.7±3.43<sup>b</sup> |

**Figure 2:**
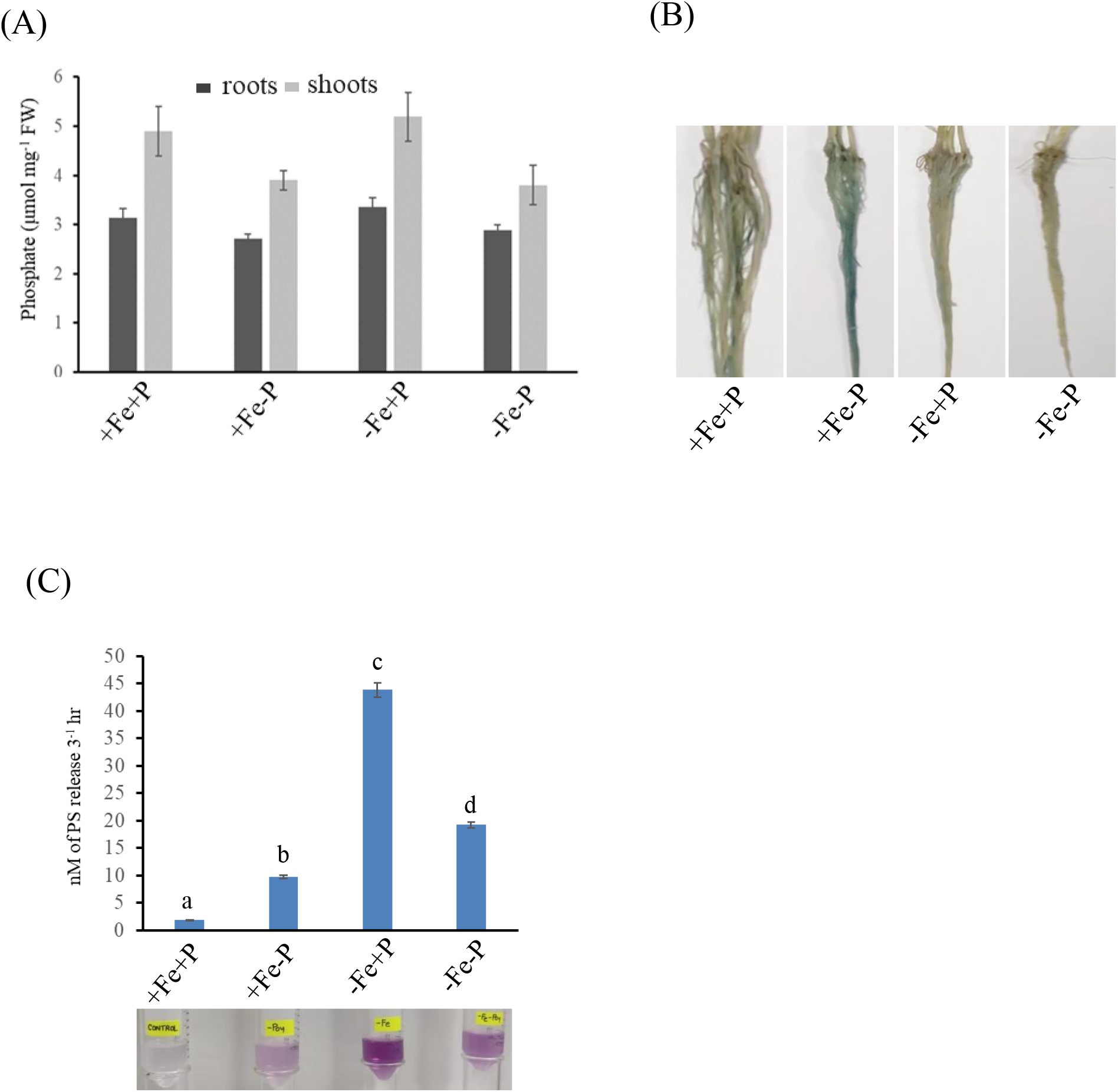
Effect of Fe and P interaction on metal accumulation and its mobilization. (A) Phosphate concentration in roots and shoots of wheat seedlings under different regimes of Fe and P, post 20 days of growth. (B) The phenotype of wheat root seedlings during the combinatorial effects of Fe and P as observed by Perl’s Stain for iron plaque (blue plaques). (C) Estimation of the phytosiderophore (PS) release (Fe-solubilization) capacity by the wheat roots under the mentioned condition. * and # indicate means are significantly different at p≤0.05 and p≤0.01, respectively (n=8).

On the other hand, wheat roots showed higher Zn accumulation under –Fe condition (Table 1). In shoots, high Fe was also significantly accumulated in –P condition relative to control. The higher accumulation of Zn was seen in shoots under –Fe+P and –Fe–P conditions, suggesting the Zn uptake is independent of P level. Interestingly, in shoots levels, Mn was high in all the treatments compared to control. Perl’s staining analysis showed no visual presence of Fe in roots under –Fe and –Fe–P conditions (Figure 2B). However, enhanced Fe-plaque colourization under –P conditions and mild staining in +Fe+P roots was observed (Figure 2B). Our data suggest that the plant roots aggressively take up Fe under –P conditions. A key aspect of Fe uptake in cereals is the release of PS for its remobilization [56]. Our assays for PS estimation revealed a high Fe-solubilizing capacity (42-47 nM) under –Fe conditions (10DAT), while this capacity decreased in –P (8-9 nM) and the lowest activity was observed in the control roots (2-3 nM; Figure 2C). Interestingly, a fairly detectable level of Fe-solubilization during –Fe–P condition was observed (16-18 nM; Figure 2C). This data indicates that P is necessary for wheat roots to respond to Fe deprivation to enhance their solubilization by releasing PS.

### 3.3 Comparative analysis of normalized RNAseq expression

The combined Fe and P deficiencies transcriptomic changes remained largely unaddressed in wheat. Therefore, to understand the molecular basis of observed phenotype and the enhanced accumulation of Fe in –P roots, we performed root RNAseq analysis in all the conditions at 20 DAT. We generated 240,013,343 quality-filtered reads, with a minimum of 89% reads having quality score >= Q30 (Table S2), for the differential expression analysis using the Kallisto-DESeq2 pipeline [37,38]. PCA plots for normalised gene expression counts revealed distinct clusters for different conditions (Figure S1A). Biological replicates for the respective four conditions (Control, –Fe, –P, –Fe–P) clustered together, thus reinforcing the reliability of the overall comparative analysis (Figure S1A and B). Refined analysis of our previous –Fe transcriptome subsequently identified 2055 up- and 2191 down-regulated genes compared to the control (Figure 3A). In response to –P condition, 2983 and 802 genes were downregulated and upregulated, respectively (Figure 3A, Table S3). Combined –Fe–P stress caused upregulation and downregulation of 1829 genes and 951 genes (Table S3). The Venn diagram revealed those unique or commonly regulated genes by Fe and P (Figure 3B, Table S4 and Table S5). Clustered heatmap analysis of all 4 transcriptomes revealed that the transcriptomes of control and –Fe–P plants were closer and distant from single Fe or P deficiency conditions (Figure S1B). Clustered heatmap analysis of the top 50 upregulated genes under combined stress across all treatments suggested that expression of these genes was similar in –Fe and – Fe–P conditions (Figure 3C, left panel). In contrast, the strongly downregulated genes (right panel) showed similar patterns during –Fe and –P. Our data suggest that wheat roots respond differently and distinctly to the combined deficiency of Fe and P.

**Figure 3:**
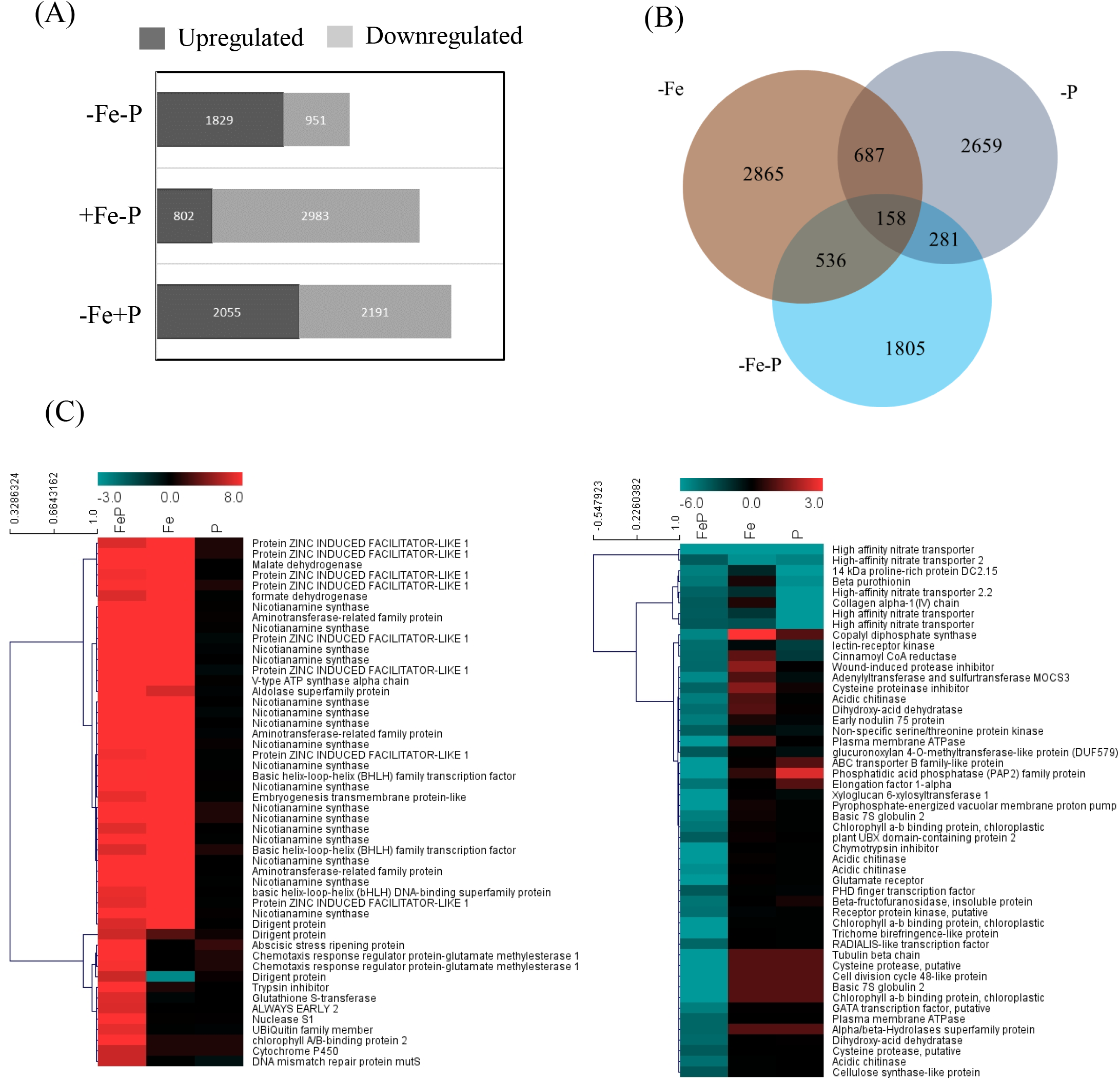
Transcriptome analysis of wheat roots grown under single (–P,–Fe) and dual (– Fe–P) conditions. (A) Genes differentially expressed in response to different deficiency conditions (Fe and P deficiency: –Fe–P; P deficiency: +Fe–P; Fe deficiency: –Fe+P) with respect to control wheat roots (+Fe+P). (B) Venn diagram represents the number of unique and common differentially regulated genes for the three conditions compared to control. (C) Clustered heatmap analysis of highest differentially responsive genes (left: genes with the highest induction; right: fifty most repressed genes) during –Fe–P condition in all three starvation conditions (combined deficiency: –Fe–P; P deficiency: –P; Fe deficiency: –Fe; Table S3). Increasing intensities of red and blue colours represent up- and downregulation (Log_2_FC), as depicted by the color scale.

### 3.4 Conserved features of response to Fe and P deficiency in hexaploid wheat

DEGs were analyzed for **–**Fe and **–**P deficiency to confirm the extent of the single deficiency. Earlier molecular signatures for the Fe deficiency response was reported [12]. Experiments for the other two deficiency conditions were also performed during the same time as the **–**Fe experiment, although sequenced and analyzed later. Since Fe deficiency response was characterized earlier [12]; we, therefore, analysed the data to identify the transcriptional changes during –P and –Fe–P. During our analysis it was observed that under **–**P, Phosphate Starvation Response (PHR1) targets such as *PHO1;H1* (TraesCS7A02G231200, TraesCS7D02G231300), *TaIPS1*(TraesCS4B02G312200), Myb TF (TraesCS2A02G100600, TraesCS2B02G117800, TraesCS2D02G100100), phosphatase-related genes like *PHOSPHATE STARVATION INDUCED 2* (PS2) orthologs (TraesCS3B02G322100, TraesCS3D02G287300) and transcripts encoding for multiple SPX containing domain proteins (TraesCSU02G094800,TraesCS7A02G554800,TraesCS7B02G478000,TraesCS7D02G541100 ,TraesCS7A02G376200,TraesCS7B02G277700 and TraesCS7D02G372600) were significantly upregulated (Table S3). This analysis confirmed a typical P deficiency response in wheat roots, as reported earlier [24]. Also, during –P, genes encoding for MADS-box TFs were highly induced (Table S6). We further evaluated phosphate starvation response (PSR) regulation during Fe and P cross-talk by analysing the overlapping expression of PSR genes (Figure 4A, TableS6). The percentage of genes overlapped between –P and –Fe –P was very low (15.79%). A total of 39 genes were commonly upregulated specifically by –P and –Fe–P, including important PSR genes like phosphate starvation-induced gene 2 (PS2). Interestingly, genes encoding for glycine-rich cell wall structural proteins were also highly upregulated in both conditions (Table S8). In contrast, 110 genes were commonly downregulated during these two conditions. Genes encoding peroxidases and proteases were notably repressed. There were 206 genes with contrasting expression in –P and –Fe–P. These included genes encoding for several germin-like protein, GDSL esterase, CytP450s, Glutathione S-Transferases, ABC transporters, *TaYS1A* and *TaVTL5* (Table S8).

**Figure 4:**
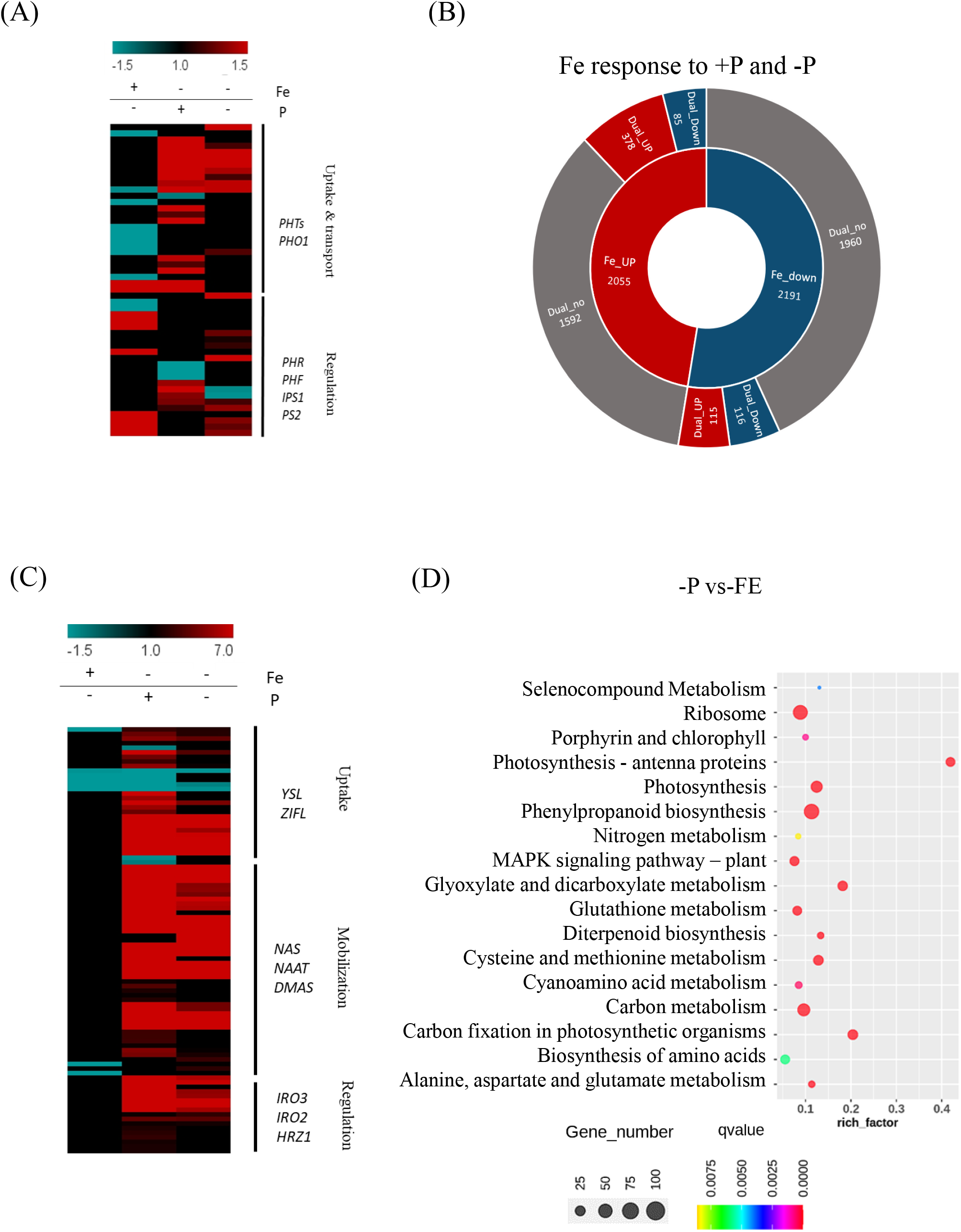
Fe and P deficiencyresponsive transcripts affected by different regimes of P and Fe. (A) Response of phosphate starvation responsive (PSR) genes to respective conditions. In total 50 PSR genes related to Pi uptake, transport and regulation were shortlisted for the analysis. (B) Effect of additional P deficiency on Fe responsive genes (Table 1). The inner circle of the sunburst graph represents transcripts up-and downregulated under Fe stress, while the outer concentric circle represents the distribution of said these genes upon dual combined stress. (UP: upregulated; Down: Downregulated; Dual_no: not differentially expressed under – Fe–P). (C) Heatmap depicting the response of 93 iron-responsive DEGs involved in Fe uptake, mobilization and regulation as identified earlier [13] during the three deficiency conditions. Red and blue bins represent up-regulation and down-regulation as represented by the scale. (D)KEGG pathway enrichment analysis for P deficiency responses w.r.t. Fe deficient wheat root transcriptome. Significantly enriched pathways (qvalue< 0.05) for +Fe–P w.r.t. -Fe+P. x-axis depicts the rich-factor, i.e., the ratio of perturbed genes in a pathway with respect to the total number of genes involved in the respective pathway, the y-axis represents the enriched pathway names, bubble sizes depict the number of genes altered in respective pathways, and increasing intensity of blue color represents increasing significance (decreasing q-value).

Similarly, the overlapping expression response of transcripts during –Fe and –Fe–P was found to be comparatively higher (24.96%; Table 3). Categorically, –Fe and –Fe–P transcriptome analysis revealed that 83.65% DEGs from –Fe alone was no longer differentially expressed under the combined –Fe–P treatment (Figure 4B, Table 3). Our analysis suggests that wheat roots could reprogram metabolic pathways in response to multiple nutrient deficiencies. In total, 494 genes were co-regulated irrespective of the presence or absence of P (Table S9). Out of these, 84 genes were observed to be significantly altered in all three deficiency conditions, including a few WRKY TFs and SPX domain-containing transcripts. There were 62 genes downregulated, including transcripts encoding for ABC-G family transporter and *TaYSL12* (Table S9). Further, 410 genes were found to be commonly regulated in Fe deficiency and dual deficiency. Those upregulated by Fe deficiency included nicotianamine synthase genes-NAS, *bHLH* and *WRKY* transcription factors, and transporters viz., *ZIFL* (zinc induced facilitator-like), *YSL, NRT1*. Genes repressed by Fe deficiency included *ABC-G* family, peroxidases, arabinogalactan protein-encoding, sulfate transporter, ferritin and loricrin-like genes. In total 200 genes depicted opposite expression patterns, depending on the presence or absence of P. These included genes encoding for no-apical-meristem (NAC)-domain-, cobalt-ion protein-encoding genes, glycosyltransferases and zinc transporter (Table S9). These genes also had co-suppressed in the –P and –Fe–P conditions but were induced under –Fe compared to the control root. These were MLO-like protein-coding genes involved in plant defence and stress response [56], sulfotransferase gene, genes encoding for lipid transfer protein, and WAT1-related protein. Furthermore, during –Fe or in –Fe–P, multiple genes involved in Fe homeostasis were also induced that suggested presence of P does not influence the expression of Fe-homeostasis related transcripts. Specifically, 93 <u>Fe s</u>tarvation responsive genes (FSR) were analyzed for their expression (Table S7). FSR genes mostly showed downregulation or no change in expression during P deficiency but were upregulated either in –Fe+P or –Fe–P (Figure 4C). These upregulated genes are known to be involved in the biosynthesis of PS *via* the methionine salvage pathway.

**Table 2:** Effect of P deficiency on Fe responsive genes. Table shows the response to Fe starvation responsive genes under an additional starvation of P (–Fe–P). (Up: upregulated; Down: Downregulated; no change: no significant differential expression under Fe–P condition).

| (–Fe) |  |  |  |
| --- | --- | --- | --- |
| (+P) |  | (–P) |  |
| Up | 2055 | Up | 378 |
|  |  | Down | 85 |
|  |  | no change | 1592 |
| Down | 2191 | Up | 115 |
|  |  | Down | 116 |
|  |  | no change | 1960 |

**Table 3:** Percentage distribution of top twenty frequent putative cis-regulatory elements (freq-pCREs) analyzed in the RNAseq data (–Fe+P, +Fe–P, –Fe–P) and in shortlisted genes involved during Fe-starvation response (FSR-93 genes) and phosphate starvation response (PSR-50 genes). Detailed list of freq-pCREs analysis is shown in Table S13.

| pCRE motifs | Percentage distribution of freq-pCREs in given condition |  |  |  |  | Top TF family |
| --- | --- | --- | --- | --- | --- | --- |
|  | –Fe+P | +Fe–P | –Fe–P | FSR | PSR |  |
| CATGCA | 63.82 | 62.22 | 65.79 | **79.57** | 80.00 | ABI3VP1 |
| ATATAT | 65.87 | 66.42 | 67.52 | **68.82** | 70.00 | ARID |
| GCATGA | 46.09 | 44.41 | 46.91 | **68.82** | 60.00 | B3 |
| ATGCAT | 64.20 | 64.68 | 65.72 | **65.59** | 70.00 | B3 |
| AATATG | 49.27 | 49.64 | 48.02 | **54.84** | 52.00 | G2-like |
| AATTAA | 54.52 | 55.30 | 54.21 | **54.84** | 56.00 | Homeobox |
| AAAGTA | 42.53 | 43.28 | 41.33 | **53.76** | 32.00 | C2C2-Dof |
| **AAACTA** | 49.29 | 50.49 | 50.65 | **51.61** | 66.00 | ARID |
| **AGCATG** | 44.21 | 44.65 | 45.07 | **51.61** | 52.00 | B3 |
| **TATGCA** | 45.24 | 46.50 | 47.05 | **50.54** | 50.00 | ABI3VP1 |
| **AGGCAT** | 33.58 | 34.35 | 33.71 | **48.39** | 38.00 | G2-like |
| **CCGGCC** | 50.82 | 49.48 | 50.40 | **48.39** | 58.00 | GeBP |
| **ACTAGT** | 43.24 | 41.80 | 41.73 | **47.31** | 32.00 | bHLH |
| **CACACG** | 43.24 | 42.03 | 42.95 | **47.31** | 28.00 | BZR |
| **CGTGCC** | 35.68 | 33.08 | 34.78 | **46.24** | 30.00 | bHLH |
| **TGTACA** | 38.86 | 40.18 | 40.18 | **46.24** | 44.00 | NAC |
| **ACGGCA** | 31.09 | 28.51 | 32.12 | **45.16** | 36.00 | NAC |
| **AGTCAA** | 38.93 | 38.39 | 40.00 | **45.16** | 58.00 | WRKY |
| **TATAGA** | 34.24 | 35.51 | 33.49 | **45.16** | 28.00 | Orphan |
| **GGCATGA** | 12.46 | 11.73 | 12.01 | **44.09** | 8.00 | B3 |

Upon comparative analysis for the single deficiencies (+Fe–P dataset compared to – Fe+P), a total of 3213 genes (660 up- and 2613 down-regulated) were differentially expressed (Table S10). Pathway enrichment analysis suggested the high representation for the phenylpropanoid pathway (PPP), carbon metabolism, carbon fixation and photosynthesis and ribosome related components under P deficiency upon comparison to –Fe (Figure 4D). While ribosome related pathway was enriched among upregulated genes, others mentioned metabolic pathways were downregulated. Multiple MADS-box TF were highly enriched in –P in comparison to –Fe. Others highly induced were ribosomal proteins, dirigent protein, Histone H2A. As expected, genes with significantly high downregulated in –P compared to –Fe include Fe-homeostasis related genes. This suggests the enrichment of distinct genes contributes to a single or combinatorial deficiency.

### 3.5. Unique signatures for –Fe–P deficiency response

Next, the comparison was drawn to identify the unique features of –Fe–P compared to control as well as single deficiency (–Fe or –P). Genes encoding Glutathione S-transferases, NBS-LRRs, chaperone-related genes, ABC and ion transporters were also highly expressed in response to –Fe– P. Additionally, three putative nitrate transporters-NRT (TraesCS5A02G388000; TraesCS7A02G428500; TraesCS3B02G285900) and a gene encoding for nitrate reductase were remarkably downregulated. These expression responses marked the characteristic of –Fe–P response in wheat roots and the downregulation of stress-responsive genes, including hydrolase and ATP binding proteins (Table S3). Specifically, eight genes encoding for UDP-glycosyltransferases were induced during –Fe–P (Table S3). A high number of phytohormone-related genes involved in auxin response and biosynthesis, including PIN and IAA sub-family genes, were significantly expressed in –Fe–P (Figure S2A & S2B; Table S11). Mapman analysis also supports the high expression of genes involved in hormone biosynthesis and secondary metabolism genes (Figure S2A & S3). Multiple TFs encoding genes forAPETALA2/ethylene response element-binding proteins (AP2/EREBPs), bHLH, MYB, WRKY and C2H2 zinc finger show high expression response in both –Fe and –Fe–P (Table S6, Table S3).

To provide insight into the distinct molecular changes caused by the additional deficiency of P in wheat roots, we studied the expression response in –Fe–P compared to –Fe. In total, 5582 genes showed an altered expression (3325 up- and 2257 down-regulated) in the dual deficiency condition when compared to the Fe stressed condition (Table S10). Among the DEGs for this analysis, most prominent were nitrate transporters (7up, 13down), GST (86up, 8down), auxin-related (22up, 20down), ABA-related (8up, 3down), a sulfur deficiency-induced 2 gene, sulfate transporters (6up, 1down), >30 ABC family transporters, ammonium transporters (6up), glycosyltransferases (63up, 23down) and ferritin (3up). Specifically, the transporters highly induced in dual deficiency compared to single Fe deficiency included nitrate and ammonium transporters, ABC transporters, nucleobase ascorbate transporters (NAT). Three homoeologs for the auxin efflux carrier family genes were induced (TraesCS5D02G361600, TraesCS5B02G356700, TraesCS5A02G354300) among other auxin homeostasis genes. Other transporters induced include phosphate transporters (TraesCS3A02G190800, TraesCS5B02G512100, TraesCS5D02G513000, TraesCS4A02G359900, TraesCS5B02G512000), *TaYSL12* homeologs, *TaZIFL7.1* (TraesCS4D02G125800, TraesCS4B02G131500). As many as 66 genes/homoeologs from the ABC subfamily were upregulated. While the highly repressed genes include encoding for terpene-synthase family gene, glutamate receptor (was also induced; 2-fold), and transporters like zinc and nitrate transporters and a VIT homoeolog (TraesCS2B02G455300). Other transcript encoding for the transporters *TaYSL9, TaYS1A, TaYSL1, TaYSL2, TaVIT2* (TraesCS5B02G202100) and *TaVIT5* (TraesCS2D02G432000, TraesCS2D02G588000), TaZIFL4.2 (TraesCS4A02G187500) were downregulated. Transcript for the glutamate/malate translocator homolog (TraesCS5D02G256100), 24 aquaporin family genes, NAT and phosphate transporters were also downregulated.

We noted that as many as 10 NAS transcripts were downregulated in the –Fe–P dual starvation compared to –Fe. The comparative analysis across conditions suggests that although NAS genes’ expression is high in both –Fe and –Fe–P compared to control, the intensity of induction is much higher in the former case. In addition, 4 *NAAT* (TraesCS1A02G291200, TraesCS1B02G300600, TraesCS1D02G289800, TraesCS7A02G334900), TaDMAS1-A (TraesCS4A02G074800, TraesCS4D02G232200) were downregulated, while a DMAS-like gene (TraesCS2A02G315600, TraesCS2B02G333900, TraesCS2D02G313700) was upregulated in – Fe–P when compared to–Fe. *AHA* (TraesCS7B02G421700) and many *IRT* transcripts were downregulated, while expression of numerous *PEZ* genes was also altered. Two genes encoding for *OsHRZ* orthologs were downregulated (TraesCS1A02G374800, TraesCS1B02G395000, TraesCS1D02G382000, TraesCS3A02G262700, TraesCS3D02G262600).

In contrast, gene enrichment analysis of –Fe–P compared to –P shows differential expression of 9290 genes (6010 up- and 3280 down-regulated; Table S10). A larger dataset was differentially expressed in –P when compared to –Fe–P. These DEGs show KEGG enrichment for metabolic pathways, including secondary metabolism and ribosome. Fe uptake and PS biosynthesis genes like *ZIFL, YSL, NAS, NAAT* were highly induced in dual starvation compared to –P, supporting Fe deficiency response during dual deficiency; however, the severity was less compared to –Fe. These analyses suggest that although the –Fe–P share few features of single deficiencies yet; multiple distinct molecular rearrangement occurs to support the physiological changes in wheat seedlings.

### 3.6 KEGG pathway enrichment analysis of DEGs and metabolome analysis

Mapman-based analysis suggests genes involved in UDP-glycosyl-transferases and Glutathione-S-transferases (GST-glutathione) related pathways during –Fe–P were more highly expressed when compared to other treatments and controls (Figure S4 and Figure 5A). Overall, our data indicate that auxin biosynthesis genes and secondary metabolism genes for lignification were highly active in the –Fe–P response (Figure S2, S3 and Figure S5) relative to other treatments. To further categorise the DEGs from each nutritional growth condition, we mapped them to the KEGG database and represented the enriched pathways by means of bubble graphs. Our analysis revealed that in response to P deficiency, genes related to the phenylpropanoid pathway (PPP), photosynthesis, ABC transporters, and genes for nitrogen metabolism were highly enriched (Figure S5). In response to –Fe–P condition, enrichment of genes involved in glutathione metabolism, glycerophospholipid metabolism, starch and sucrose metabolism and galactose metabolism pathways and cysteine and methionine metabolism was observed (Figure 5B). Interestingly, enrichment of cysteine and methionine metabolism genes was also observed in response to –Fe+P treatment [12], indicating that the enrichment of these genes is Fe specific and independent of P status. Next, we analyzed the significantly altered pathways in the dual deficiency (–Fe–P) compared to –Fe and –P datasets to highlight the unique and common pathway elements among these conditions (Figure 5B). KEGG analysis for the enriched pathways in the –Fe–P treatment compared to –Fe indicates that Cysteine and methionine metabolism, Phenylalanine, tyrosine and tryptophan biosynthesis, sulfur metabolism, flavonoid biosynthesis, Vitamin B6 metabolism, Diterpenoid biosynthesis were highly enriched in –Fe–P treatment compared to–Fe. While upon extracting the pathways specifically enriched in –Fe–P relative to–P were for ribosome synthesis, glyoxylate and dicarboxylate metabolism, inositol phosphate metabolism, photosynthesis, porphyrin and chlorophyll metabolism, phosphatidylinositol signalling system, and ABC transporters were highly represented. This suggests a distinct pathway restructuring during the changing regimes of P and Fe.

**Figure 5:**
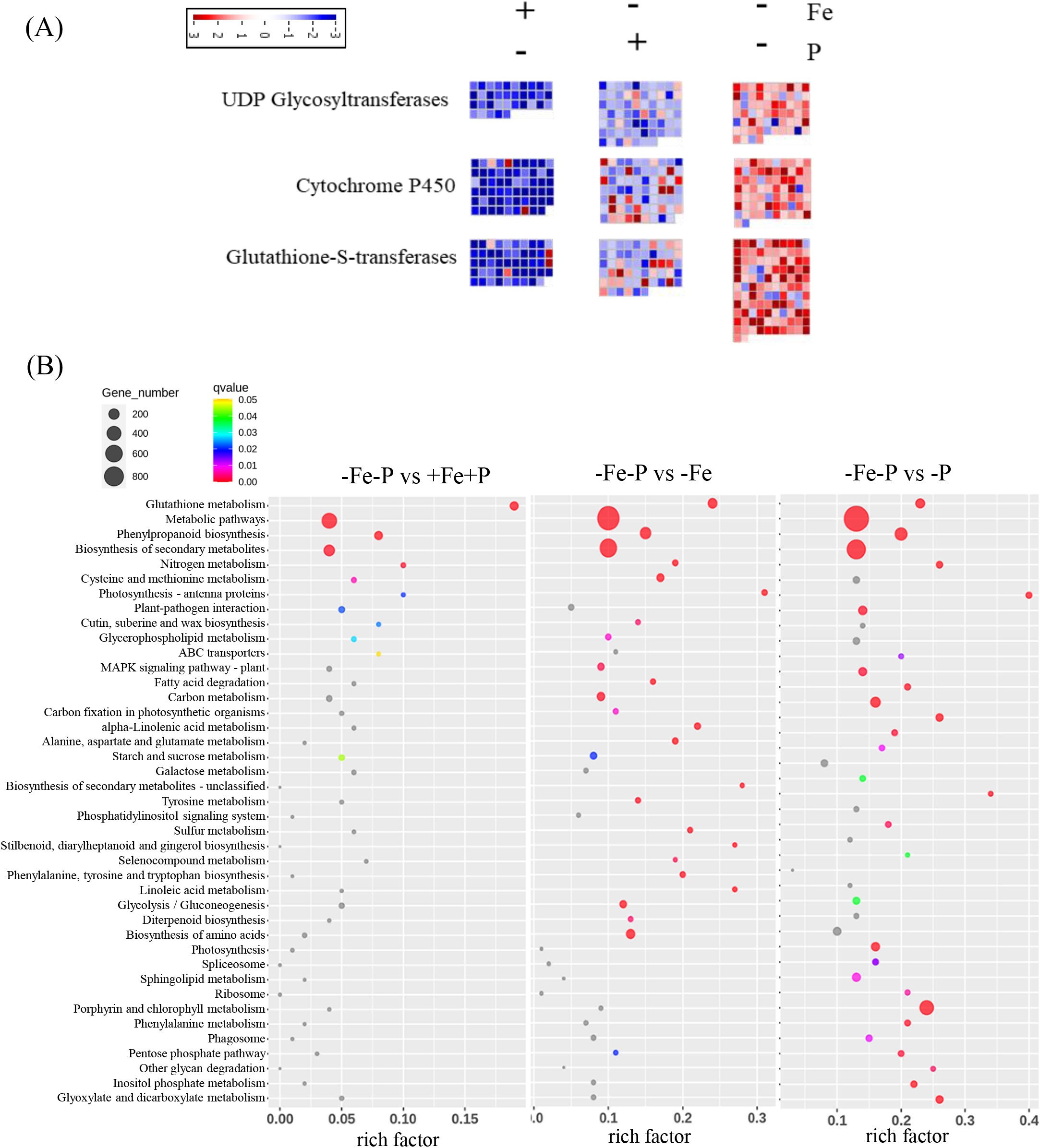
Mapman and KEGG Pathway enrichment analysis. (A) Detailed analysis of DEGs for UDP glycosyltransferases, cytochrome P450 and Glutathione-S-transferases. Log2 fold change values of the DEGs were imported into MapMan. Red and blue bins represent up-regulation and down-regulation as shown by the scale. (B) Significantly enriched pathways (qvalue< 0.05) for –Fe–P (combined deficiency) compared to +Fe+P (control). Significantly enriched pathways (qvalue< 0.05) for –Fe–P (combined deficiency) compared to –Fe+P (Fe deficiency). Significantly enriched pathways (qvalue< 0.05) for –Fe –P (combined deficiency) and +Fe–P (P deficiency). x-axis depicts the rich-factor, i.e., the ratio of perturbed genes in a pathway with respect to the total number of genes involved in the respective pathway, the y-axis represents the enriched pathway names, bubble sizes depict the number of genes altered in respective pathways and increasing intensity of the color represents increasing significance (q-value).

Next, we studied the role of primary metabolites, using the GC-MS analysis. Our analysis showed significant variation in the accumulation of metabolites during the nutrient deficiency treatments (Figure 6A, Table S12). While suppression of oxalic acid and increase in 4-ketoglucose levels was unique for –P treatment, the increase in fumaric acid and myo-inositol marked specific metabolic change for –Fe conditions. A contrasting level of serine and succinic acid in +Fe–P (low) and –Fe+P (high) was found to be normalized in the combined deficiency relative to the control. The –Fe–P conditions is characterized by specific metabolic changes marked by a decrease in acetic acid, butanoic acid, valine, threonine and glucofuranoside levels and increased accumulation of β-amino-isobutyric acid, strearic acid, arabinonic acid and aconitic acid. Nevertheless, few metabolites decreased in both the –Fe–P and –Fe conditions such as aspartic acid, hexonic acid, glucose cystathione and, alanine. While the acids showed high accumulation in –Fe, they were highly reduced during –Fe–P. The metabolites that decrease in –Fe–P appeared to follow the same trends in –P conditions, including citric acid and hexapyranose. Finally, sugars, sugar conjugates (i.e., d-ribofuranose, a-d-galactopyranoside, a-d-mannopyranoside), amino acids (i.e., b-aminoisobutyric acid, cystathione and L-alanine), aconitic acid and arabinonic were predominant in –Fe–P conditions (Figure 6B). Overall, our analysis linked the accumulation of specific metabolites that were distinct for a particular condition.

**Figure 6:**
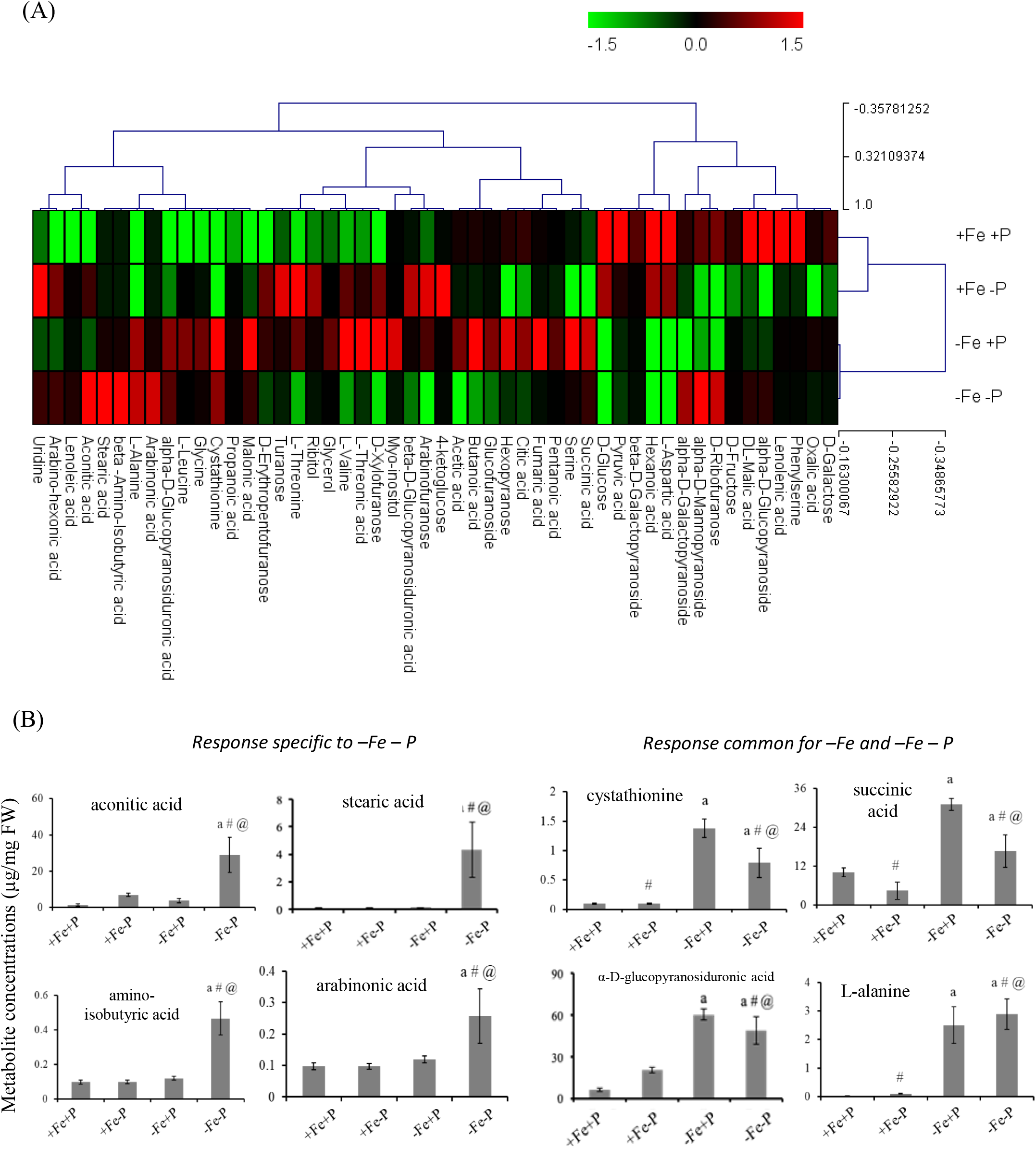
Overview of the changes in metabolome in roots of wheat seedlings subjected to different growth regimes of Fe and P. (A) GC-MS based heatmap comparison of root metabolites in +Fe +P; -Fe +P; +Fe –P and –Fe –P condition. Metabolites are sorted according to their classes, specifically, sugars, general acids, amino acids, sugar conjugates, fatty acids and polyols. Concentrations (µg/mg, fresh weight) of individual metabolites in test conditions were compared with control metabolite concentrations and represented with Log_2_FC values as shown by the scale. Data means ±SD of n=3 experiments. (B) Quantitative plot for the metabolite concentrations (μg/mg FW) for response specific to –Fe –P and response common in –Fe and –Fe–P. Different symbols indicate significant differences between the conditions determined by Fisher’s LSD (p < 0.05). +Fe +P, control; -Fe +P, Fe deficiency; +Fe –P, P deficiency; –Fe –P, Fe and P deficiency. ‘a’ represents a significant difference against control, ‘#’ represents significant difference against Fe deficiency and ‘@’ represents significant difference against P deficiency.

### 3.7 Enrichment of Fe responsive cis-regulatory elements in DEGs

Fe-responsive cis-regulatory elements were analyzed in the promoter of differentially expressed genes during –Fe, –P, and –Fe–P conditions to get insight into transcriptional regulation. Our analysis revealed multiple TFs that are predominantly expressed in the –Fe–P condition compared to P deficiency (Table S6, Figure S6). Genes encoding for multiple bHLH, C2H2 and NAC TFs were highly represented in –Fe–P. Previously, a comprehensive resource of new putative frequent cis-regulatory elements (freq-pCREs), was also identified in the gene clusters responding to Fe deficiency [14,57]. Herein, we checked for the predominance of freq-pCREsin the gene-promoters that were differentially regulated in response to single (–Fe and – P) or combined –Fe–P deficiencies using in-silico methods. Although, this study was done using the wheat Ensembl (Chinese Spring) database, yet recent genome sequencing of wheat cv. C-306 [35] was used to confirm the promoter survey observation of this study. This homology matches were confirmed by performing the blast analysis of ten randomly selected FSR gene promoter sequences from Chinese Spring that resulted in the >99% identity for each of these genes in C-306. Since, the strategy-II mode of Fe and P uptake remains largely conserved among the wheat cultivars, we extended our study to FSR and PSR related genes from the wheat Ensembl database.

Further, to optimize and validate our analysis, we used three control sets of genes (∼100) with Log_2_FC between -0.5 and 0.5 in –Fe with respect to control and checked for the occurrence of freq-pCREs in their promoters. Uniform representation of freq-pCREs was detected in triplicates across each set, thus validating our assessment (Table S13). We then tested this module on a subset of FSR genes and PSR genes. Interestingly, when FSR and PSR-related genes were analyzed, this balance was found to be perturbed. The dominance and biasness of freq-pCREs were observed significantly in Fe deficiency, as represented in Table 4. For example, cis-domains such as GGCATGA, CACGTC, AAAGTA, ACTAGT, CACACG, AATTGC and CGTGCC were present in higher percentages in Fe deficiency responsive genes as compared to P deficiency. Similarly, cis-elements for phosphate responsive regions such as P1BS (PHR1 binding sequences, GNATATNC) were present at 62% of the PSR and 12.90% of FSR genes (Table S14). Interestingly, the P1BS motif was present 34.97, 34,61% and 36.91% in the promoter regions of DEGs in response to –Fe+P, +Fe–P and –Fe–P, respectively. Our work revealed an overlapping response of DEGs during –Fe–P deficiency as freq-pCREs were highly enriched in Fe deficiency and present in the promoters of P-response related genes.

## 4. Discussions

### 4.1 Transcriptome profiling uncovers the link between Fe and P interactions

Fe and P are essential elements for plants, utilized in nearly every cellular process. However, there is limited understanding of Fe and P homeostasis in crops to coordinate the physiological and molecular responses. The current work aimed to fill this knowledge gap by providing insights into the wheat responses to Fe and/or P deficiency stresses that could help cope with varying soil-environmental conditions. Our data showed that P deficiency compensates for the negative effect of Fe deficiency on wheat growth and development. Our transcriptome analysis suggests the enriched presence of putative Fe responsive cis-regulatory binding sites. Combining transcriptome and metabolome analysis revealed a specific component underlying the combined Fe and P deficiency stress response in wheat.

The components of the plant response to P starvation, including the PHR1-controlled IPS1-miR399-PHO2 signaling cascade, remains conserved across the plant kingdom, including wheat [24,58]. In our –P wheat root transcriptome analysis, the negative regulator, *SPX1* orthologs, was upregulated in accordance with earlier studies noted in rice and wheat [24,59].*TaIPS*1 induction was complemented by *PHO1* and multiple *PHT* genes being downregulated. However, upregulation of *PHO1;H1* rice orthologs was observed. *RNS1* and *RNS3*were upregulated in earlier P-starvation studies in rice and wheat but downregulated in our case, which could be a potential marker for the late response to P-starvation. Wheat ortholog for *AtTIR1* (Transport Inhibitor Response 1), an auxin receptor that is involved in specifically lateral root elongation under –P [60] was found to be induced (TraesCS1D02G099900). Other genes like *LPR1, LPR2, PDR2, STOP1* and *ALMT1*, which are central to root architectural changes, specifically reduction in primary root growth, in *Arabidopsis*, showed no altered expression in –P condition. However, *ALMT* and *STOP1*-like genes show altered expression in –Fe conditions irrespective of P presence (Table S3).Also, SCARECROW (SCR), a TF involved in regulating root patterning and stem cell maintenance under –P [61] was not differentially expressed under –P, but altered in the –Fe transcriptome (TraesCS4B02G141400, TraesCS7B02G077800), while 2 SCR genes (TraesCS3A02G360300, TraesCS3B02G392600, TraesCS3D02G354100, TraesCS7B02G078300, TraesCS7D02G174100, TraesCS7D02G174300) were upregulated in –Fe–P.

To survive in Fe or P-deficient soils, crop plants like rice, maize, and soybeans have adapted multiple strategy responses. However, crop plant responses to co-occurring Fe and P stresses remained poorly understood. We observed that common genes expressed in response to –Fe and –Fe–P supports the Strategy-II mode for Fe uptake even during changing regimes of P. For instance, the relatively high release of PS and gene expression patterns of certain specific YSL, metal transporters and a few TFs involved in metal homeostasis either in –Fe or –Fe–P confirms that wheat primarily utilizes Strategy-II mode of Fe uptake route to mobilize Fe under P deficiency (Figure S7) [62]. Overall, relative high gene expression surge for the FSR genes was observed in Fe deficiency responses, compared to –Fe–P. It was interesting to observe the release of PS even during –P, which could be attributed to the expression of multiple wheat TOM genes [63]. Although the expression of wheat TOM genes was lowered during –P compared to –Fe, yet the PS release during –P points to some alternate mechanism for Fe mobilization. Interestingly, accumulation and release of PS in barley roots were shown to be dependent on the P levels [64]. Based on this, we conclude that, under –Fe condition, optimal P is necessary for enhanced PS accumulation and release. Conversely, ABCG transporters have been speculated to release phenolics and flavins in *Arabidopsis* to mobilize Fe to roots [65]. However, such substrate analysis for wheat ABC transporter for PS secretion ability remains to be tested. Another possibility is that under –P, multiple organics acids (citrate, malate) are known to be accumulated in wheat roots, enhancing the Fe-solubilization [12]. In future, it will be, therefore, interesting to investigate how –P could cause the release of PS during this condition.

Although –P condition led to the increase in the root Fe, most of the Strategy II and alternate Fe uptake genes like IRT, NRAMP, ABC transporter genes are either unaltered or downregulated. Strategy II genes like TaYS1A and DMA Synthesis genes are downregulated in –P as observed in rice [52], while these are upregulated in –Fe–P. Interestingly, *TaVTL5* was found to be upregulated in –P and repressed in both –Fe conditions. This could be to avoid the Fe toxicity due to increased Fe concentration in –P roots. Genes involved in root to shoot phosphate transfer were induced under –Fe condition, explaining the increased P concentration in –Fe shoot. IDEF1 regulates –Fe responses in graminaceous plants. Although IDEF1 core-binding site (CATGC) containing freq-pCREs, <u>CATGC</u>C and C<u>CATGC</u>were present in FSR gene promoters, yet their occurrence was more predominant in PSR genes. This might suggest a probable IDEF1 role in regulating the increased expression of PSR genes in Fe deficiency. Previously, phylogeny analysis was done across the wheat genomes that suggested close proximity (>97.2) of C-306 with Chinese spring cv [35]. Our extended analysis of the cis-element in the strategy-II genes in the Chinese spring (Wheat Ensembl) seems to be conserved across the wheat genomes thereby implying the conserved role of Fe and P homeostasis across hexaploid wheat genomes. Further, such genetic variations need to be dissected on case-by-case basis.

Holistic rearrangement of metabolites under Fe, P and other nutrient deficiencies are necessary to cope with the stress conditions [53]. Our study revealed few specific signatures for the combinatorial deficit (–Fe–P) (Figure 6B). Our data further confirmed that under P and Fe deficiencies,plant rootaccumulates organic acids, including malate and citrate, for efficient mobilization of nutrients [66–70]. The accumulation of multiple organic acids is also well reflected in our study during the combined stress condition. Under P deficiency, wheat decreases citrate levels (Table S12 and Figure 6A). Fe deficient wheat roots tend to accumulate high citrate levels with downregulation of citrate synthase transcript (TraesCS7A02G409800, TraesCS7D02G403000). Citrate being a Fe(III) chelator has been reported to play a relevant role in iron acquisition and xylem Fe transport [71,72]. Overexpression of citrate synthase has been shown to assist plant growth during–P by increasing citrate exudation [73]. Our transcriptome changes support these metabolic changes. Therefore, it could be speculated that the release of the citrate under –P could support efficient mobilization of Fe. Lastly, accumulation of other metabolites, including sugars, sugar conjugates, organic acids, amino acids, fatty acids and polyols, was also observed. Earlier, similar observations for accumulating these metabolites were reported in barley and wheat [74,75]. Altogether, our metabolome analysis suggests strategic rearrangement in wheat roots to ensure P availability and Fe mobilization. Such cross-talk between P and Fe to regulate Fe uptake and transport has also been reported in dicots, such as *Arabidopsis*. For instance, P deficiency was shown to induce Fe homeostasis-related genes like *AtFRO2, AtIRT1* and ferritin genes (*AtFer1)* [18,76,77]. Likewise, Fe deficiency can induce the expression of a subset of genes involved in the P acquisition genes [43,78]. These data, along with our observations, points to the unique perspective of plant P and Fe interactions.

### 4.2 Core components of dual deficiency response

Identifying the specific signatures for the combined deficiency of –Fe and –P will provide an important link for their homeostatic interactions. This study led to the identification of particular signatures at the transcript and metabolome levels. In plants, the PPP is the primary source of numerous phenylalanine derivatives involved in multiple developments and physiological processes, including lignin biosynthesis and cell wall development [79,80]. The role of glycosyltransferases has been demonstrated to control the phenylpropanoid pathway genes [81] efficiently. The high expression of UGT transcripts during –Fe–P suggests reorganization of metabolic pathways that resulted in the defining of molecular signatures. Our MapMan enrichment analysis supports the high expression of genes especially encoding for simple phenols, lignin biosynthesis, isoflavonoids and carotenoids (Figure S5). The high cell wall-related activity in wheat roots could be correlated with enhanced root biomass allocation (Figure 1E). Previously it was observed that lignin biosynthesis could be linked to excess Fe-related responses to provide tolerance in rice [81]. The affirmative role of lignification and the cytochrome P450 in wheat needs functional attention to address their roles under combined deficiencies.

KEGG enrichment analysis reinforced our finding that genes encoding for glutathione metabolism were significantly enriched during –Fe–P treatment. Levels of Glutathione and its metabolism have been correlated with the tolerance for –Fe [12,82,83]. The role of glutathione under metal stress is well documented [84,85]. In our study, the role of glutathione is reflected by the accumulation of its precursor cystathionine under –Fe as well as in –Fe–P (Figure 6). This suggests that glutathione expression is proportional to the Fe stress in roots. Therefore, one could postulate that, high expression of GST coupled with the glutathione accumulation could be one of the hallmarks Fe deficiency ‘gateway modules’ that prevent roots against oxidative damage. Other components, such as nitric oxide-mediated iron uptake response, are controlled by glutathione supply [85]. The robust expression of glutathione-related genes in – Fe–P condition suggests that roots undergo the redox process required for survival. Accumulation of glycine and serine has been implicated to negatively affect root length and nitrate uptake in *Brassica campestris* [86].Our data coupled with the previous studies indicated that contrasting glycine and serine accumulation levels in –Fe conditions could be accounted for short root development [12]. Glycine can induce ethylene-guided inhibition of root elongation, or it could be converted into aminobutyric acid moieties during stress [86,87].

The growth during Fe deficiency is in contrast to the decreased total biomass under P deficiency, and this effect was also reflected in the combined deficiency, i.e., Fe–P. However, this phenomenon was different in the *Arabidopsis*, where the primary roots were shorter under –P and recovered by –P–Fe [88]. This indicates that dicots and monocots have evolved distinct genetic programs to respond to single or combined nutrient stress. In contrast, *Arabidopsis* roots show a reduction in the primary root length in +Fe–P [89], whereas in wheat, the effect is reversed with enhanced root growth in –P. Thus, based on our analysis, P-deficient wheat roots could enhance nutrient (Fe, Zn and Mn) accumulation, reprogram primary metabolism (oxalate and citrate) and affect the combination of PSR and FSR responses including those involved in auxin homeostasis irrespective of the plant species [30,89].

Plants respond to nutrient stress by shuffling the biomass allocation [90]. It was also reported that due to nutrient stress, levels of sugars/carbohydrates might be perturbed. In this study, the root to shoot biomass allocation was perturbed during the single and combined stress suggesting a conserved mechanism among the plants. Our GC-MS data suggests increasing levels of different sugars (Figure S8) in –Fe condition compared to other stresses. This indicates an increase in the carbohydrate metabolism-related activities to enhancing the biomass allocation in shoots. On similar lines, sucrose accumulates in *Arabidopsis* roots under –Fe condition and promotes auxin signalling [91]. These observations in *Arabidopsis* and wheat are consistent to account for the changes in biomass allocation in stress conditions. The deficiency of individual nutrients like Fe or P affects wheat and other crops for the overall physiological growth and development; and yield components [92,93]. In wheat, we observed short root under –Fe condition, the rescue of root phenotype by simultaneous –Fe–P stress due to changes in biomass allocation. The enhanced GST dependent auxin activity could explain this compensation effect that influences the number of lateral roots and root adaptation for the combined deprivation stress. GST-mediated regulation of auxin pathway for root architecture re-arrangement has been previously reported in plants [94].

## 5. Conclusions

Our work highlights the molecular and biochemical changes that are responsible for readjustments for the dual deficiency conditions. Based on the observation, we propose a model for the interaction of Fe and P at the rhizospheric level (Figure S9). The effect of P and Fe deficiency although distinct could provide an interplay between FSR and PSR genes. Early studies in *Arabidopsis* suggested a working model for the root growth in P deficiency stress contrasts with those observed in rice and wheat despite a consistent accumulation of Fe under P limitation. In *Arabidopsis*, the overaccumulation of Fe in –P is considered the leading cause of the inhibition of root growth [31]. But in wheat, we showed that the overaccumulation does not inhibit root growth in –P conditions. We propose that the overaccumulation of Fe in roots of P-deficient wheat could be accounted for by the persistent release of PS and solubilization by organic acids in the rhizosphere. Our work opens new research avenues to uncover the molecular basis of P and Fe signalling cross-talk in plants. It will help design strategies to develop wheat cultivars with better Fe and P use efficiency and deficiency tolerance.

## Acknowledgements

The authors thank Executive Director, NABI, for facilities and support. This work was funded by the NABI-CORE grant to AKP. GK and VS acknowledge NABI-SRF Fellowships. Technical help from Jagdeep Singh for the GC-MS analysis is highly appreciated. DBT-eLibrary Consortium (DeLCON) is acknowledged for providing timely support and access to e-resources for this work. We appreciate editing for the English language by Dr Mandy Kendrick, Missouri State University, USA. The wheat genome resources developed by International Wheat Genome Sequencing Consortium are highly appreciated.

## Author Contributions

Conceptualization, AKP and HR; methodology, GK, VS, VM, DS and PK; formal analysis, GK, AKP, VS, SM and VM; investigation, GK, VS and AK; resources, JS, HR; data curation, GK and VS; writing—original draft preparation, AKP, VS and GK; writing—review and editing, AKP, VM, PK, and GK; supervision, AKP, JS SM, HR; project administration, AKP; funding acquisition, AKP. All authors have read and agreed to the published version of the manuscript.

