## Supplementary material for "Physiological and molecular responses to combinatorial iron and phosphate deficiencies in hexaploid wheat seedlings": New Final Supp Figures S1_S9.pptx

### Slide 1
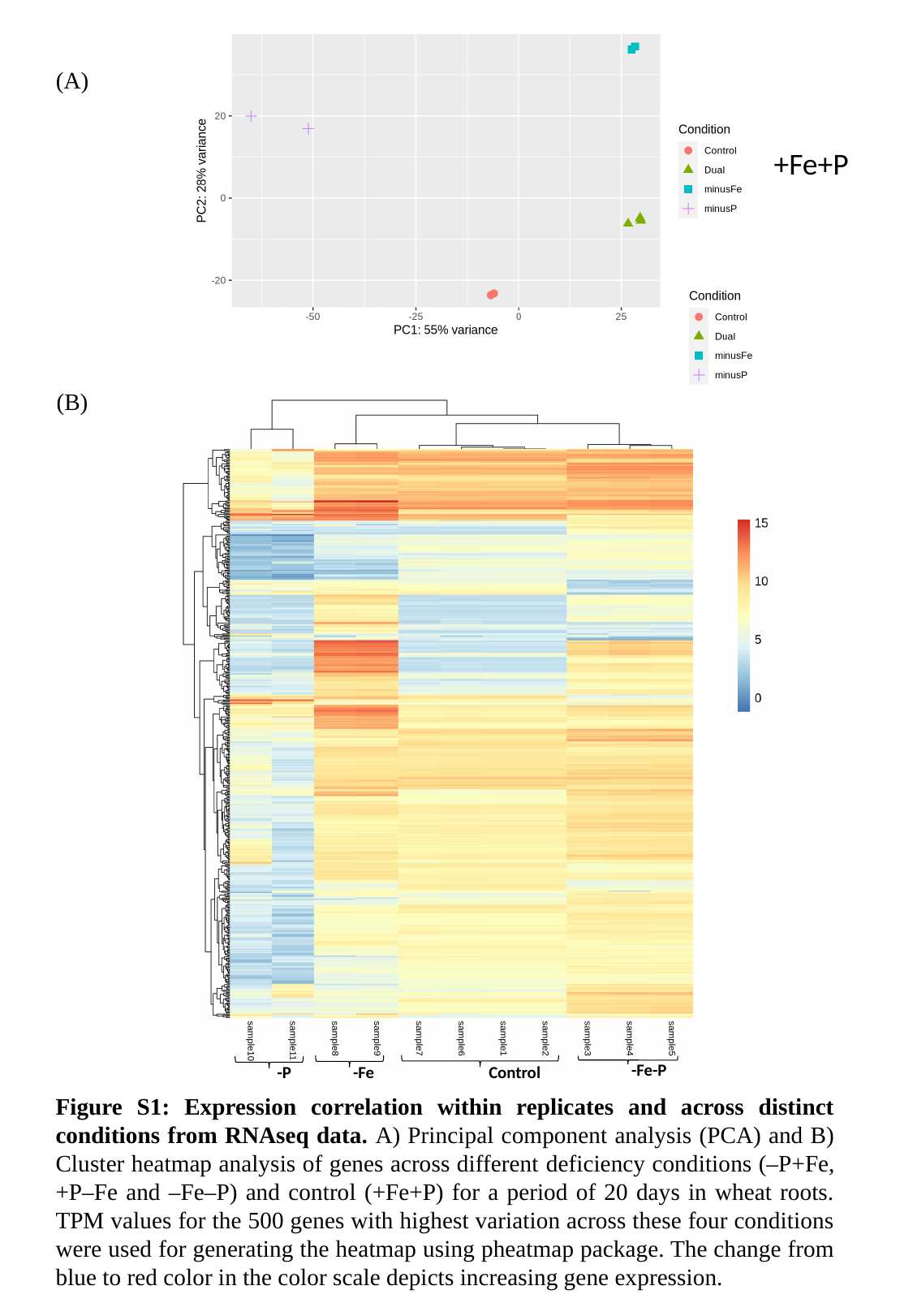

(A)
+Fe+P
(B)
-Fe-P
Control
-P
-Fe
Figure S1: Expression correlation within replicates and across distinct conditions from RNAseq data. A) Principal component analysis (PCA) and B) Cluster heatmap analysis of genes across different deficiency conditions (–P+Fe, +P–Fe and –Fe–P) and control (+Fe+P) for a period of 20 days in wheat roots. TPM values for the 500 genes with highest variation across these four conditions were used for generating the heatmap using pheatmap package. The change from blue to red color in the color scale depicts increasing gene expression.

### Slide 2
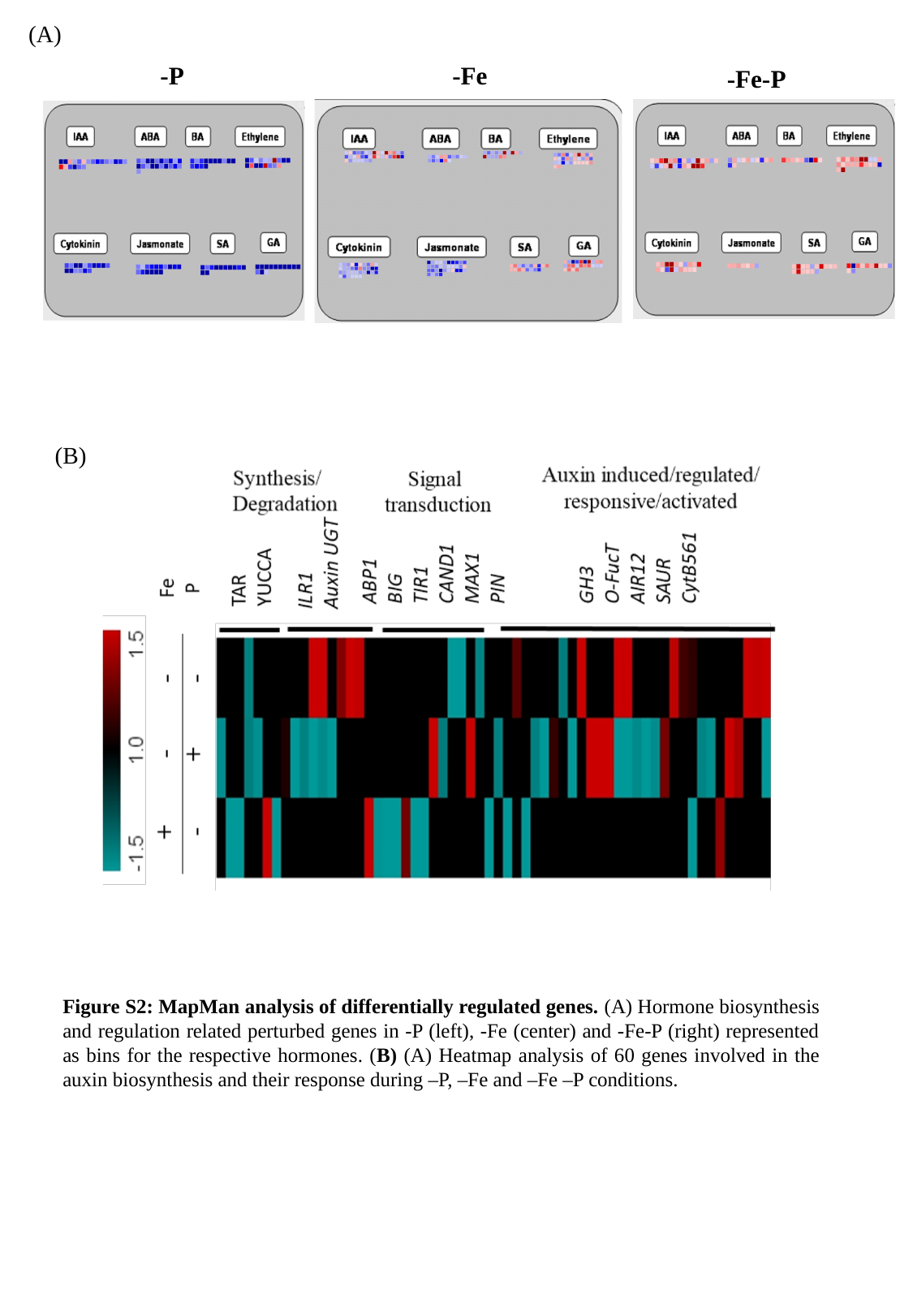

(A)
-P
-Fe
-Fe-P
(B)
Figure S2: MapMan analysis of differentially regulated genes. (A) Hormone biosynthesis and regulation related perturbed genes in -P (left), -Fe (center) and -Fe-P (right) represented as bins for the respective hormones. (B) (A) Heatmap analysis of 60 genes involved in the auxin biosynthesis and their response during –P, –Fe and –Fe –P conditions.

### Slide 3
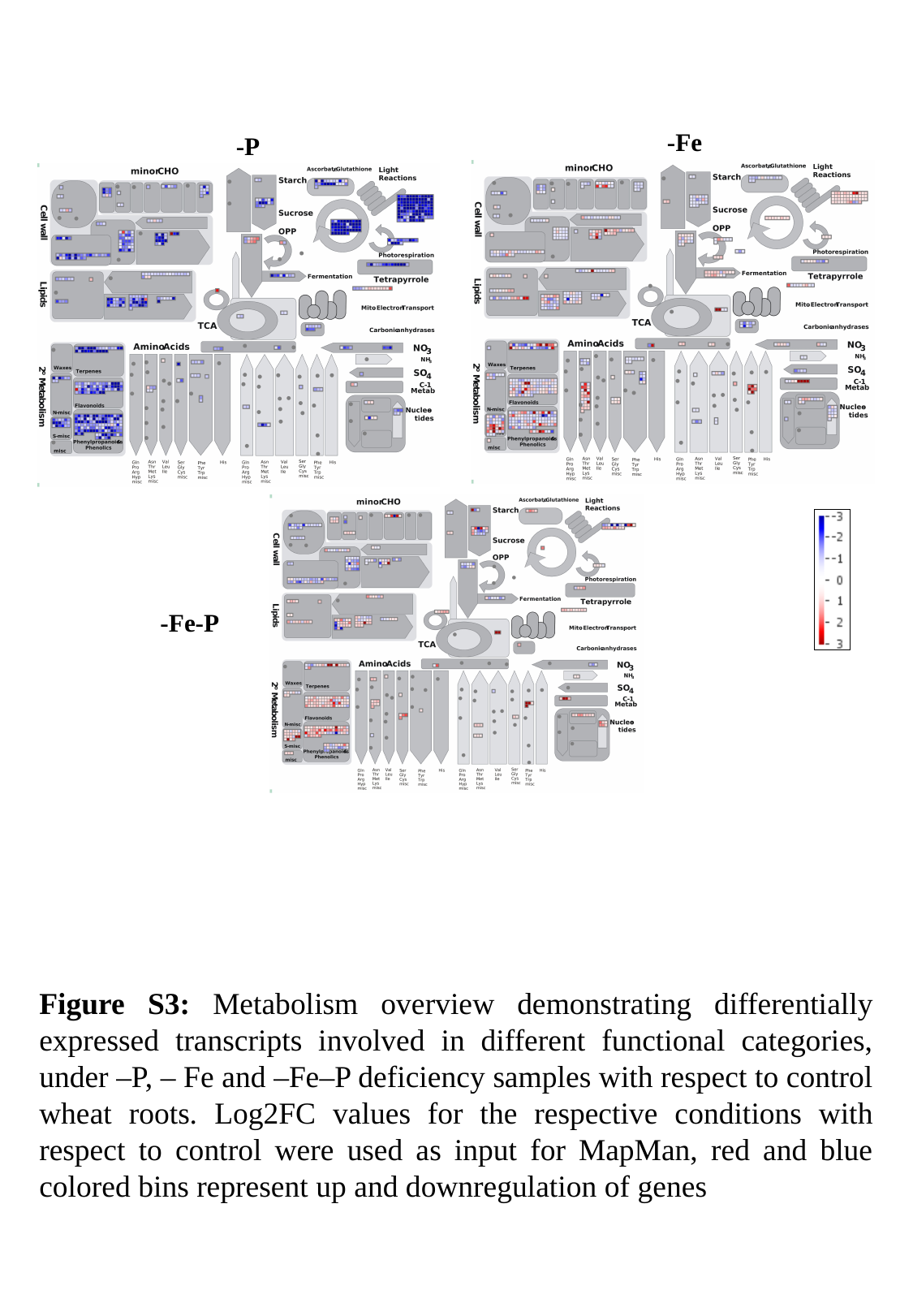

-Fe
-P
-Fe-P
Figure S3: Metabolism overview demonstrating differentially expressed transcripts involved in different functional categories, under –P, – Fe and –Fe–P deficiency samples with respect to control wheat roots. Log2FC values for the respective conditions with respect to control were used as input for MapMan, red and blue colored bins represent up and downregulation of genes

### Slide 4
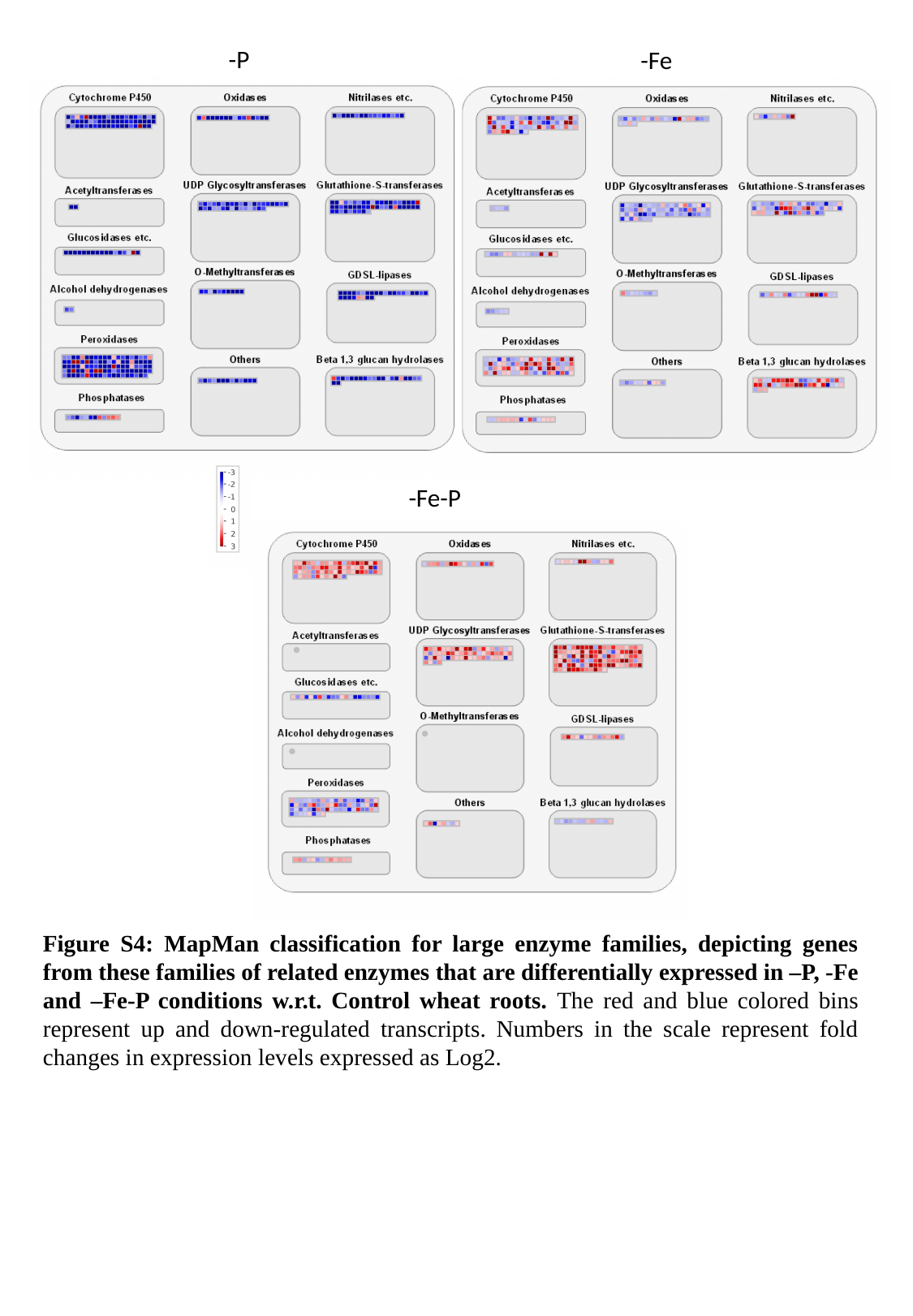

-P
-Fe
-Fe-P
Figure S4: MapMan classification for large enzyme families, depicting genes from these families of related enzymes that are differentially expressed in –P, -Fe and –Fe-P conditions w.r.t. Control wheat roots. The red and blue colored bins represent up and down-regulated transcripts. Numbers in the scale represent fold changes in expression levels expressed as Log2.

### Slide 5
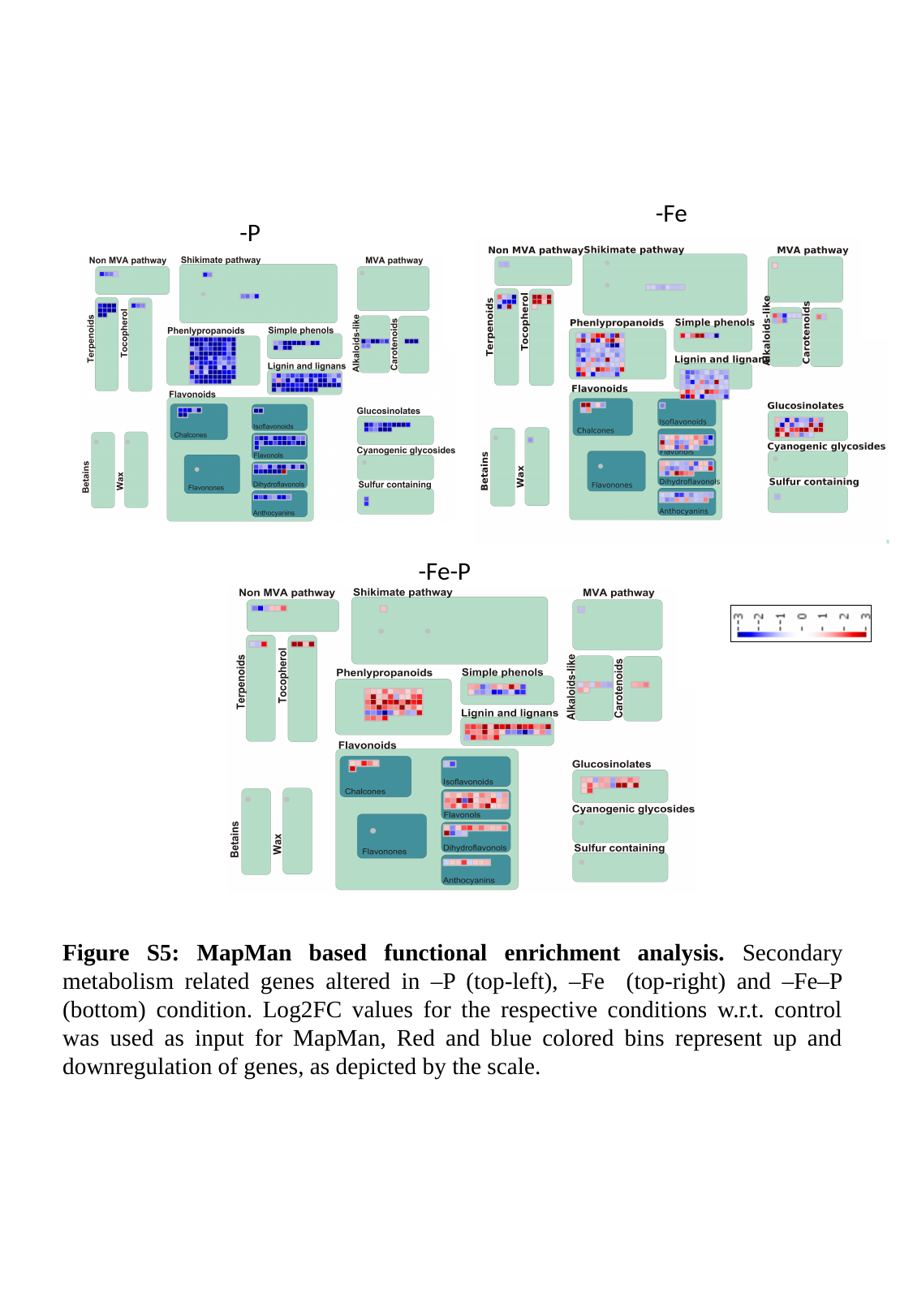

-Fe
-P
-Fe-P
Figure S5: MapMan based functional enrichment analysis. Secondary metabolism related genes altered in –P (top-left), –Fe (top-right) and –Fe–P (bottom) condition. Log2FC values for the respective conditions w.r.t. control was used as input for MapMan, Red and blue colored bins represent up and downregulation of genes, as depicted by the scale.

### Slide 6
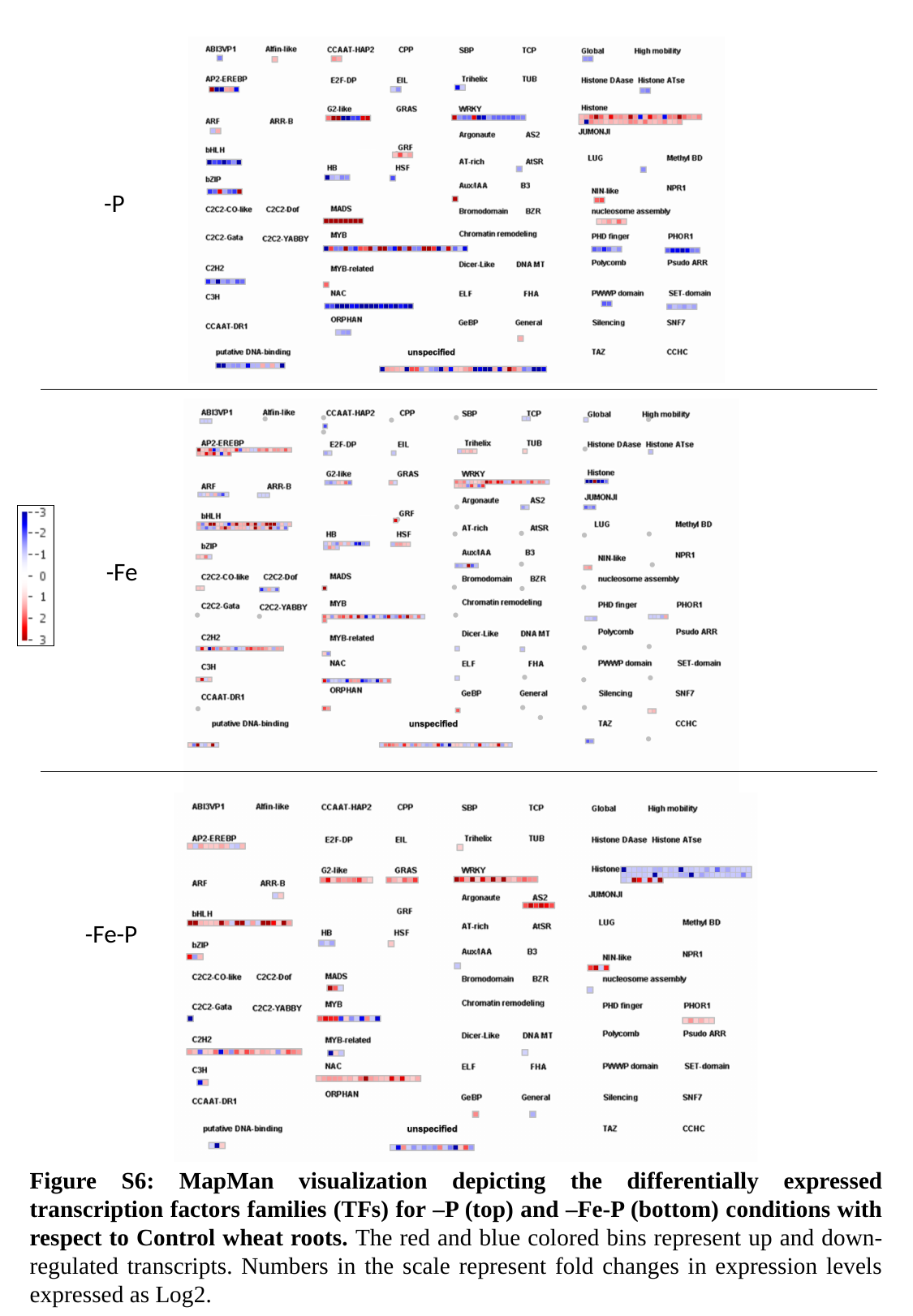

-P
-Fe
-Fe-P
Figure S6: MapMan visualization depicting the differentially expressed transcription factors families (TFs) for –P (top) and –Fe-P (bottom) conditions with respect to Control wheat roots. The red and blue colored bins represent up and down-regulated transcripts. Numbers in the scale represent fold changes in expression levels expressed as Log2.

### Slide 7
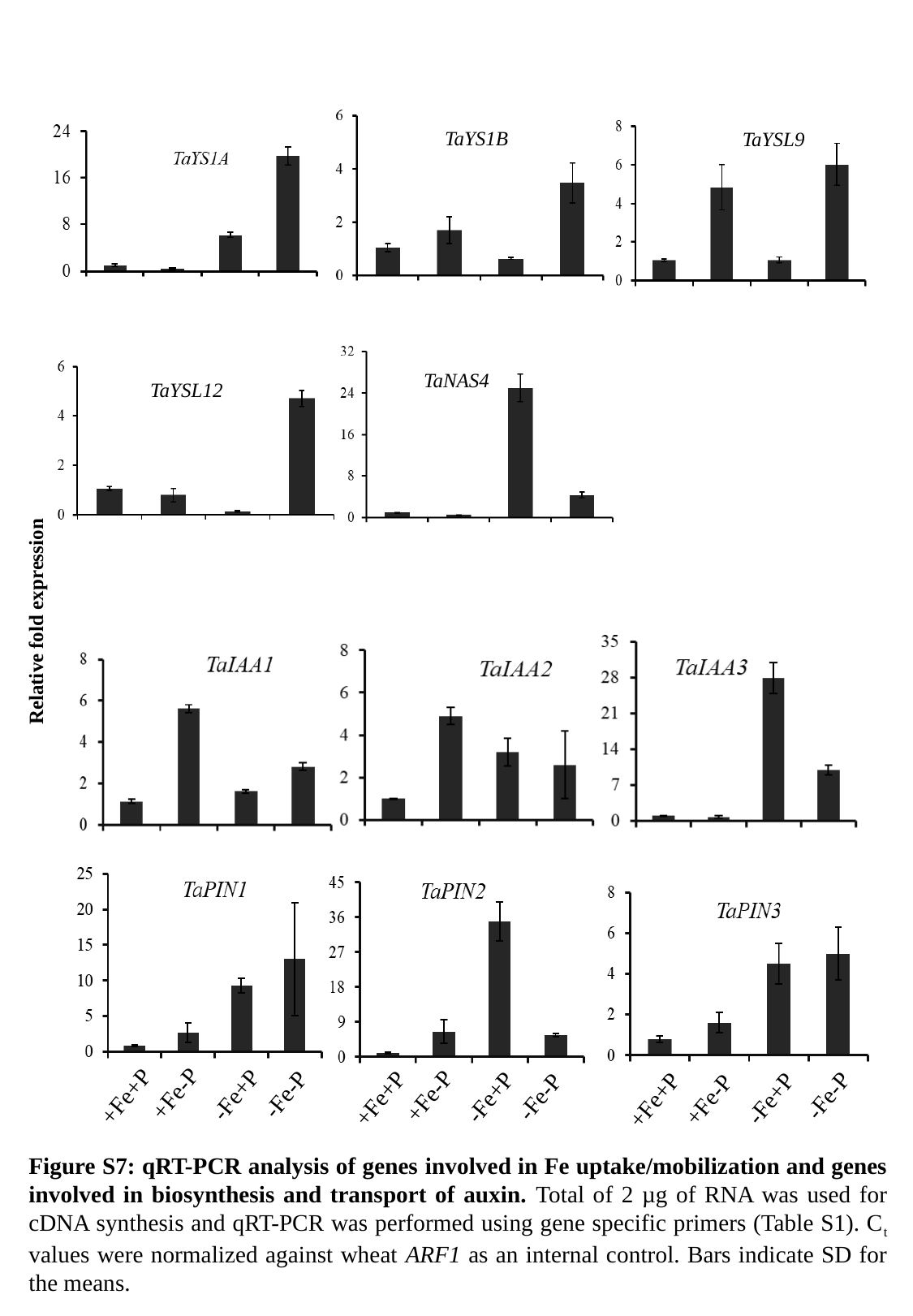

TaYS1B
TaYSL9
TaNAS4
TaYSL12
Relative fold expression
+Fe-P
-Fe-P
-Fe+P
-Fe-P
+Fe-P
+Fe+P
-Fe-P
-Fe+P
-Fe+P
+Fe+P
+Fe+P
+Fe-P
Figure S7: qRT-PCR analysis of genes involved in Fe uptake/mobilization and genes involved in biosynthesis and transport of auxin. Total of 2 µg of RNA was used for cDNA synthesis and qRT-PCR was performed using gene specific primers (Table S1). C_t_ values were normalized against wheat ARF1 as an internal control. Bars indicate SD for the means.

### Slide 8
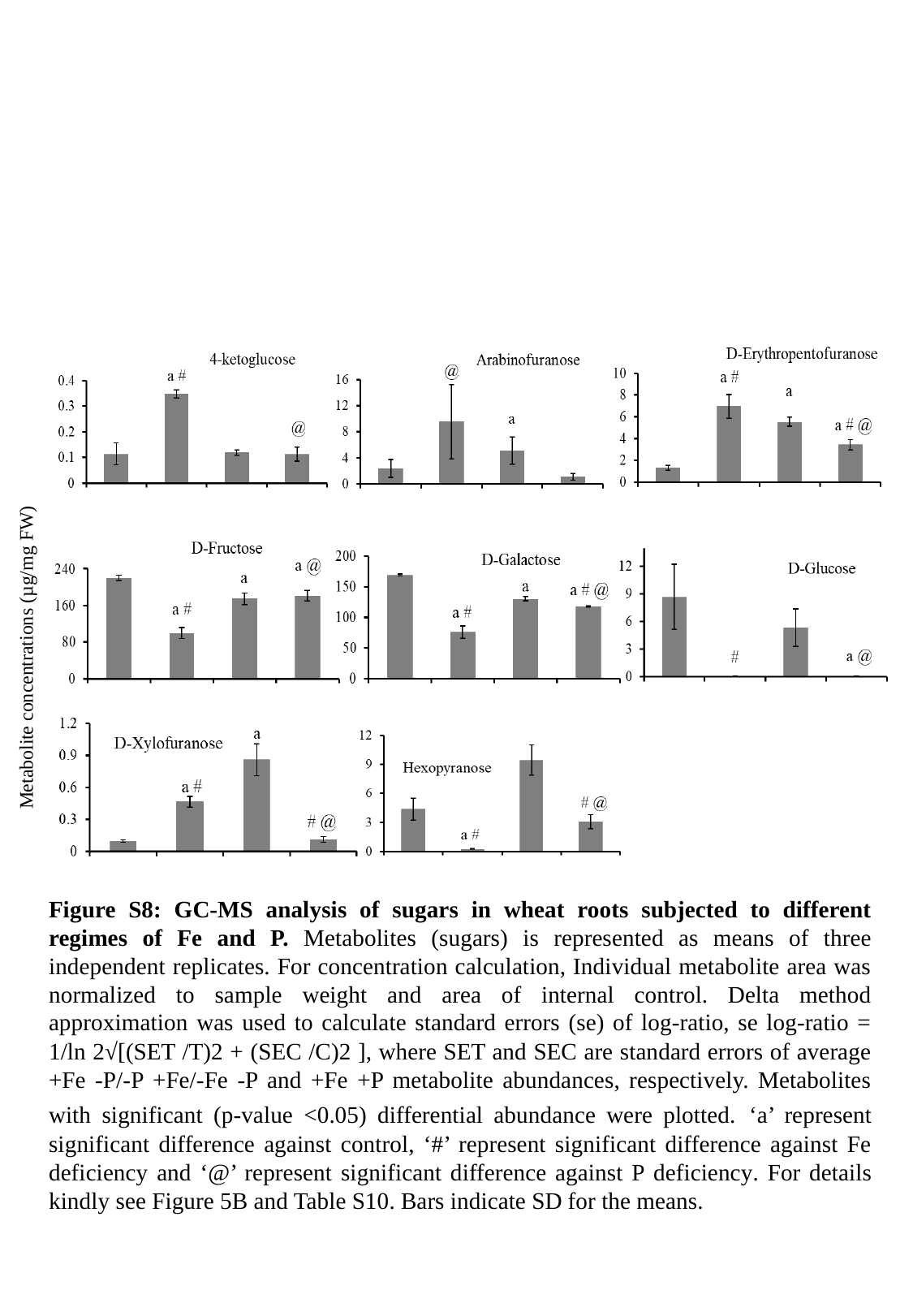

Metabolite concentrations (µg/mg FW)
Figure S8: GC-MS analysis of sugars in wheat roots subjected to different regimes of Fe and P. Metabolites (sugars) is represented as means of three independent replicates. For concentration calculation, Individual metabolite area was normalized to sample weight and area of internal control. Delta method approximation was used to calculate standard errors (se) of log-ratio, se log-ratio = 1/ln 2√[(SET /T)2 + (SEC /C)2 ], where SET and SEC are standard errors of average +Fe -P/-P +Fe/-Fe -P and +Fe +P metabolite abundances, respectively. Metabolites with significant (p-value <0.05) differential abundance were plotted. ‘a’ represent significant difference against control, ‘#’ represent significant difference against Fe deficiency and ‘@’ represent significant difference against P deficiency. For details kindly see Figure 5B and Table S10. Bars indicate SD for the means.

### Slide 9
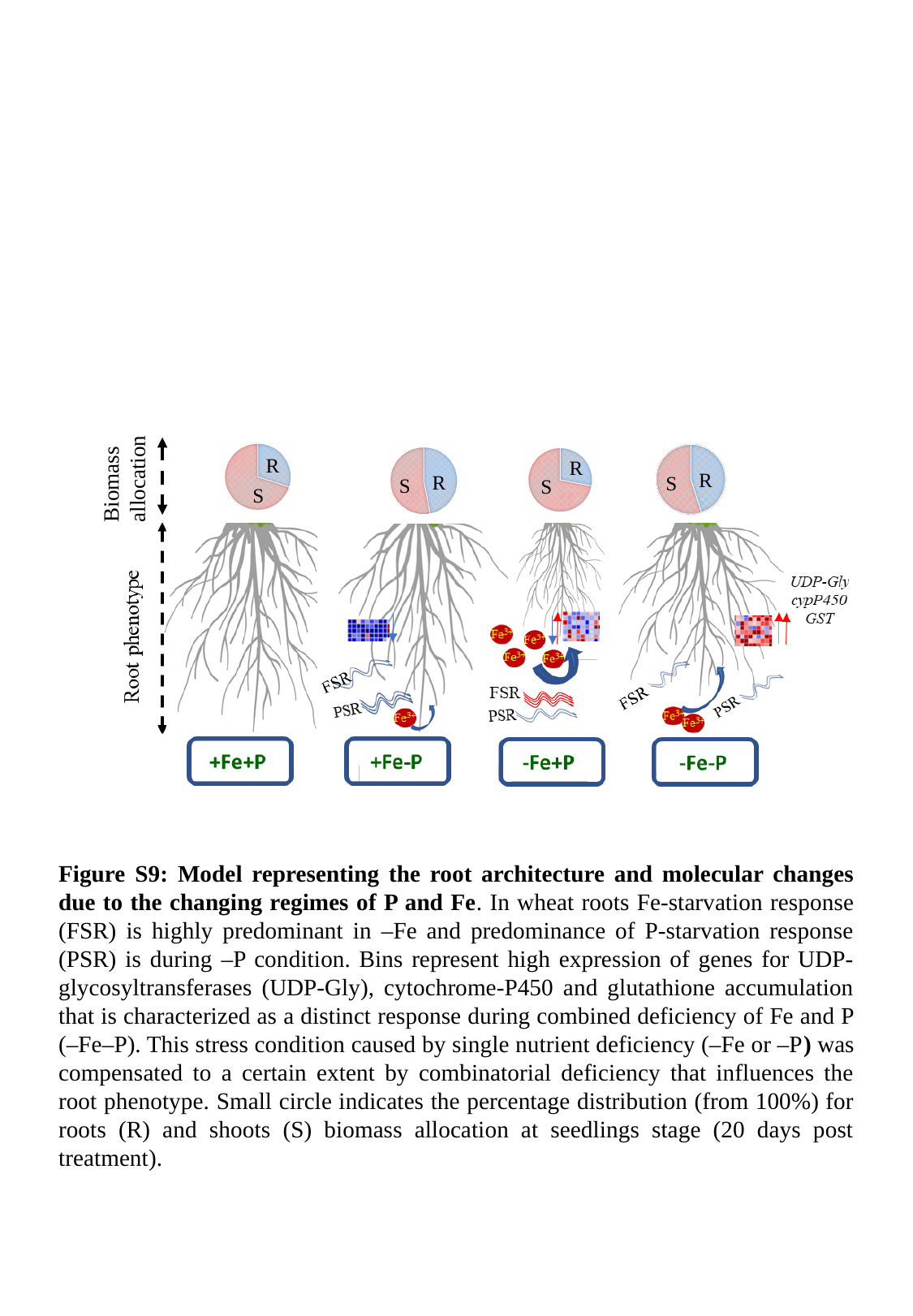

Biomass
allocation
R
R
R
R
S
S
S
S
Figure S9: Model representing the root architecture and molecular changes due to the changing regimes of P and Fe. In wheat roots Fe-starvation response (FSR) is highly predominant in –Fe and predominance of P-starvation response (PSR) is during –P condition. Bins represent high expression of genes for UDP-glycosyltransferases (UDP-Gly), cytochrome-P450 and glutathione accumulation that is characterized as a distinct response during combined deficiency of Fe and P (–Fe–P). This stress condition caused by single nutrient deficiency (–Fe or –P) was compensated to a certain extent by combinatorial deficiency that influences the root phenotype. Small circle indicates the percentage distribution (from 100%) for roots (R) and shoots (S) biomass allocation at seedlings stage (20 days post treatment).
