## Supplementary material for "Physiological and molecular responses to combinatorial iron and phosphate deficiencies in hexaploid wheat seedlings": Table S1 Primers Used.docx

**Table S1: List of primers used in the current study.** **wheat genes named according to rice RAP-DB/RefSeq based on KOBAS annotation.*

| Gene name | Primer sequence 5’-3’ | Amplicon size in bp |
| --- | --- | --- |
| *TaARF1* | F:TGATAGGGAACGTGTTGTTGAGGC | 234 |
|  | R:AGCCAGTCAAGACCCTCGTACAAC |  |
| *TaYS1A* | F:ATGACATACGTTGGTGCCGGGATGATTTGCCC | 202 |
|  | R:CCCCCATGATGAGAGCTATGCATATGAAGGCC |  |
| *TaYS1B* | F: AGCTTGCTGGACCGCTTCGGCATCGTG  R: CCAATCCCTGGCTCCTTGTAGCTCCCC | 206 |
| *TaYSL9* | F:CGGGTCCTACCTGCTCGGGCTC  R:CTGAGAGGGACAAGCGCGAGAATCC | 156 |
| *TaYSL12* | F: GAAGAAGAACAGCACCATCCCGGTCTCG  R: ATGTAGTACCACTTGAGGTCCGGGAAGATC | 207 |
| *TaZIFL4.2* | F: CATATGAAGTGTCTCCGGTAGCCATATGCA  R: AGGTCTTGTTGTCGGTCCAGCCATTGGAGG | 214 |
